# TAPER: Pinpointing errors in multiple sequence alignments despite varying rates of evolution

**DOI:** 10.1101/2020.11.30.405589

**Authors:** Chao Zhang, Yiming Zhao, Edward L Braun, Siavash Mirarab

## Abstract

Erroneous data can creep into sequence datasets for reasons ranging from contamination to annotation and alignment mistakes. These errors *can* reduce the accuracy of downstream analyses such as tree inference and *will* diminish the confidence of the community in the results even when they do not impact the analysis. As datasets keep getting larger, it has become difficult to visually check for errors, and thus, automatic error detection methods are needed more than ever before. Alignment masking methods, which are widely used, completely remove entire aligned sites. Therefore, they *may* reduce signal as much as or more than they reduce the noise. An alternative is designing targeted methods that look for errors in small species-specific stretches of the alignment by detecting outliers. Crucially, such a method should attempt to distinguish the real heterogeneity, which includes signal, from errors. This type of error filtering is surprisingly under-explored. In this paper, we introduce TAPER, an automatic algorithm that looks for small stretches of error in sequence alignments. Our results show that TAPER removes very little data yet finds much of the error and cleans up the alignments.

---

Multiple sequence alignments used in phylogenetics and other evolutionary analyses are susceptible to errors. The input to phylogenetic inference is often prepared through a long pipeline of several error-prone steps (Fig. 1). Together, these steps can leave datasets riddled with many types of errors, called *data pipeline errors* here. For example, contaminating DNA (Simion *et al.*, 2018; Laurin-Lemay *et al.*, 2012; Olson and Hassanin, 2003) and sequencing errors can persist even after the assembly of sequences (Francois *et al.*, 2020; Breitwieser *et al.*, 2019). The process for establishing homology using genome annotations, whole genome alignment, or sequence matching involves complex computational problems (Lunter *et al.*, 2008), and thus, errors in homology are not just possible but rampant (Springer and Gatesy, 2018). Most commonly used methods further assume *orthology*, and errors in orthology detection are common (Laurin-Lemay *et al.*, 2012; Salichos and Rokas, 2011). Alignment errors are also ubiquitous and can impact tree accuracy (Li-San Wang *et al.*, 2011; Liu *et al.*, 2009; Ogdenw and Rosenberg, 2006; Fletcher and Yang, 2010; Smirnov and Warnow, 2020). The prevalence of these errors in phylogenomic datasets has been appreciated (Springer and Gatesy, 2018, 2016; Hosner *et al.*, 2016; Sayyari *et al.*, 2017; Philippe *et al.*, 2017; Laurin-Lemay *et al.*, 2012), and several phylogenomics studies have now been criticized (Springer and Gatesy, 2018, 2016; Gatesy and Springer, 2014; Jeffroy *et al.*, 2006; Salichos and Rokas, 2013; Shen *et al.*, 2017). Data pipeline errors represent a major source of that criticism.

**Fig. 1.**
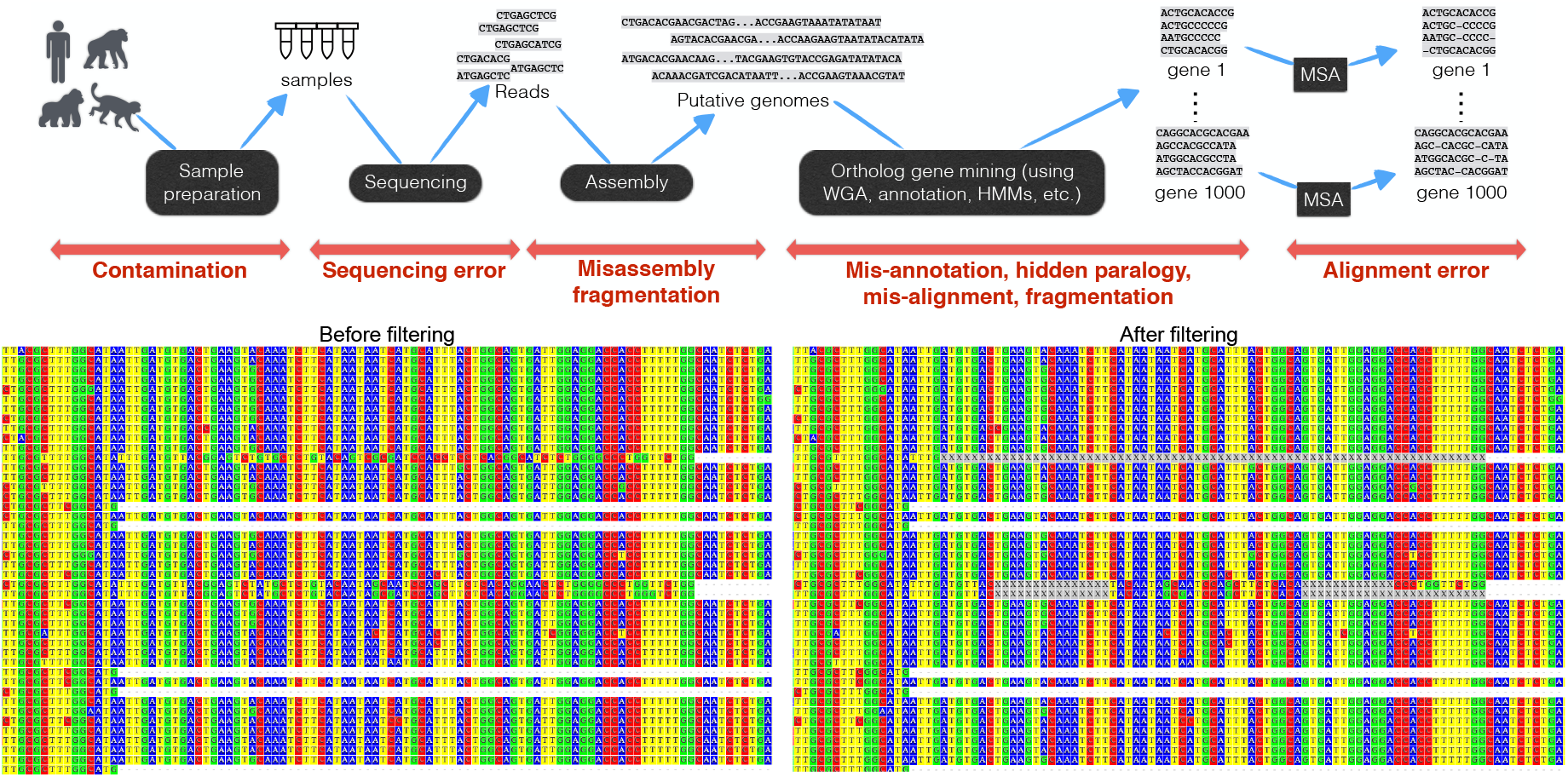
Data pipeline errors. Top: Many error-prone steps are needed to produce gene multiple sequence alignments (MSAs) used as input to phylogenomic reconstruction methods. Bottom: an example data pipeline error in the avain dataset of Jarvis *et al.* (2014), identified by Springer and Gatesy (2018), where in gene CEPT1, for three species (Phoenicopterus, Mesitornis, Leptosomus) Intron 3 is aligned with exon 4 of other taxa (left). TAPER is able to detect most but not all of the mis-aligned parts for these sequences (X positions).

Despite the widespread recognition of the challenges associated with data pipeline errors, there are no truly satisfactory ways to address those errors. Researches can not afford to visually curate their phylogenomic datasets. Many authors have mentioned the need for better methods for detecting errors in data (e.g., Philippe *et al.*, 2017). However, detecting and eliminating error comes with its own caveats and issues, and caution is warranted. Aggressive filtering of data upon any suspicion of error can add bias (e.g., by eliminating highly-divergent but genuinely homologous site patterns that should be considered in analyses). Thus, excessive filtering has the potential to eliminate signal. Consistent with this explanation, studies have found that commonly used alignment masking algorithms often have limited impact on accuracy (Portik and Wiens, 2020) and can even reduce the accuracy of phylogeny inference (Tan *et al.*, 2015). Regardless of whether small errors *actually* impact the tree inference, a question on which there is disagreement, the existence of data pipeline errors has the *potential* to impact the results, which can diminish confidence in analyses. Thus, even when errors do not impact the analyses, researchers benefit from detecting and removing them as long as removing the errors does not remove signal. Achieving this objective requires error detection methods that are targeted and find *minimal* portions of the data with putative errors.

The existing methods for data filtering mostly focus on finding entire genes or entire species that should be eliminated (e.g., Hosner *et al.*, 2016; Molloy and Warnow, 2018; Huang *et al.*, 2016). Somewhat more targeted are alignment masking methods that eliminate entire sites from a sequence alignment in order to avoid mis-alignment (e.g., Castresana, 2000; Dress *et al.*, 2008; Capella-Gutiérrez *et al.*, 2009; Rajan, 2012). Another form of filtering is to keep all genes and all species but to remove specific species from specific genes because of fragmentation (Sayyari *et al.*, 2017), evidence of unexpected patterns of tree topology and branch length (de Vienne *et al.*, 2012; Wickett *et al.*, 2014; Mai and Mirarab, 2018), or detection of rogue taxa (Westover *et al.*, 2013).

Existing trimming methods operate at coarse levels. Many forms of pipeline errors can be limited to a small stretch of a sequence in a particular species (not all species) and in a particular set of positions (not an entire gene). For example, the dominant from of error that Springer and Gatesy (2018) found in the avian data of Jarvis *et al.* (2014) relates to small pieces of introns being annotated as exons (e.g., Fig.1b). Such errors are limited to a handful of sequence in a small stretch of sites. Eliminating entire genes, entire sites, or entire species because a small stretch includes errors wastes data. We need methods to find specific stretches of a specific species in a specific gene that appear erroneous.

Finding such small stretches of errors can be formulated as outlier positions in a multiple sequence alignment (MSA). Imagine a MSA that is almost fully conserved for all species across all sites except that a small stretch from a single species is close to random with respect to other sequences aligned to it. Such outliers *can* be detected. One would hope alignment methods would leave such sequences unaligned, but most commonly used alignment methods are known to over-align (Löytynoja and Goldman, 2008; Loytynoja and Goldman, 2005). Thus, these stretches are suspect and likely to be erroneous.

Alignment trimming using the outlier detection paradigm needs to contend with two related issues. On the one hand, sequence divergence among species is a function of their phylogenetic relationships and evolutionary rates, and simply being divergent from other sequences cannot be viewed as evidence of error. If one naively looks for species that look unusually divergent compared to the remaining sequences, an outgroup or an ingroup with a highly accelerated rate of evolution would be mistakenly taken as erroneous (a false positive error detection). On the other hand, rates of evolution change across sites, and thus, how much divergence is “normal” depends on the sequence context.

We advocate for two-dimensional (2D) error detection: finding stretches in a sequence that are unusually divergent compared to other sequences, calibrating the *normal* level of divergence based on both genomic positions (columns) *and* the species (rows). An outlier should be detected only if a sequence is unusually divergent along both axes. For tree-based trimming, de Vienne *et al.* (2012) has pioneered such a two-dimensional approach in their mathematically elegant method Phylo-MCOA, and TreeShrink (Mai and Mirarab, 2018) follows a similar philosophy. However, we are not aware of two-dimensional outlier methods for sequence data. The most relevant method is DivA (Zepeda Mendoza *et al.*, 2014), which does look for stretches to remove from individual sequences using sliding windows, but is only applicable to amino acid sequences.

In this paper, we introduce the Two-dimensional Algorithm for Pinpointing ERrors (TAPER) that takes a multiple sequence alignment as input and outputs outlier sequence positions. Using both simulated and real data, we show that TAPER is able to pinpoint errors in multiple sequence alignments without removing large parts of the alignment.

## Description

### 2D outlier detection algorithm

We first describe our general-purpose 2D outlier detection algorithm (Alg. 1). The input is a set of *n* aligned sequences with length *L* on any arbitrary alphabet Γ (e.g., nucleotide) plus missing data (e.g., gaps). The output is a delineation of each sequence into alternating normal and outlier regions. We use *letter* to refer to a position in a sequence, not counting gaps (or ambiguous letters like X for amino acid) as letters.

#### Step 1

Compute a divergence score for each letter *c* in each column *i* of the alignment. Small scores should indicate agreement with a strong consensus in that site and the largest values should indicate disagreement from an otherwise strong consensus. High scoring letters are candidate outliers. Thus, the drop from large values to small values should be not be gradual; instead, it should have a fast drop as deviation from the consensus weaken.

**Algorithm 1.**
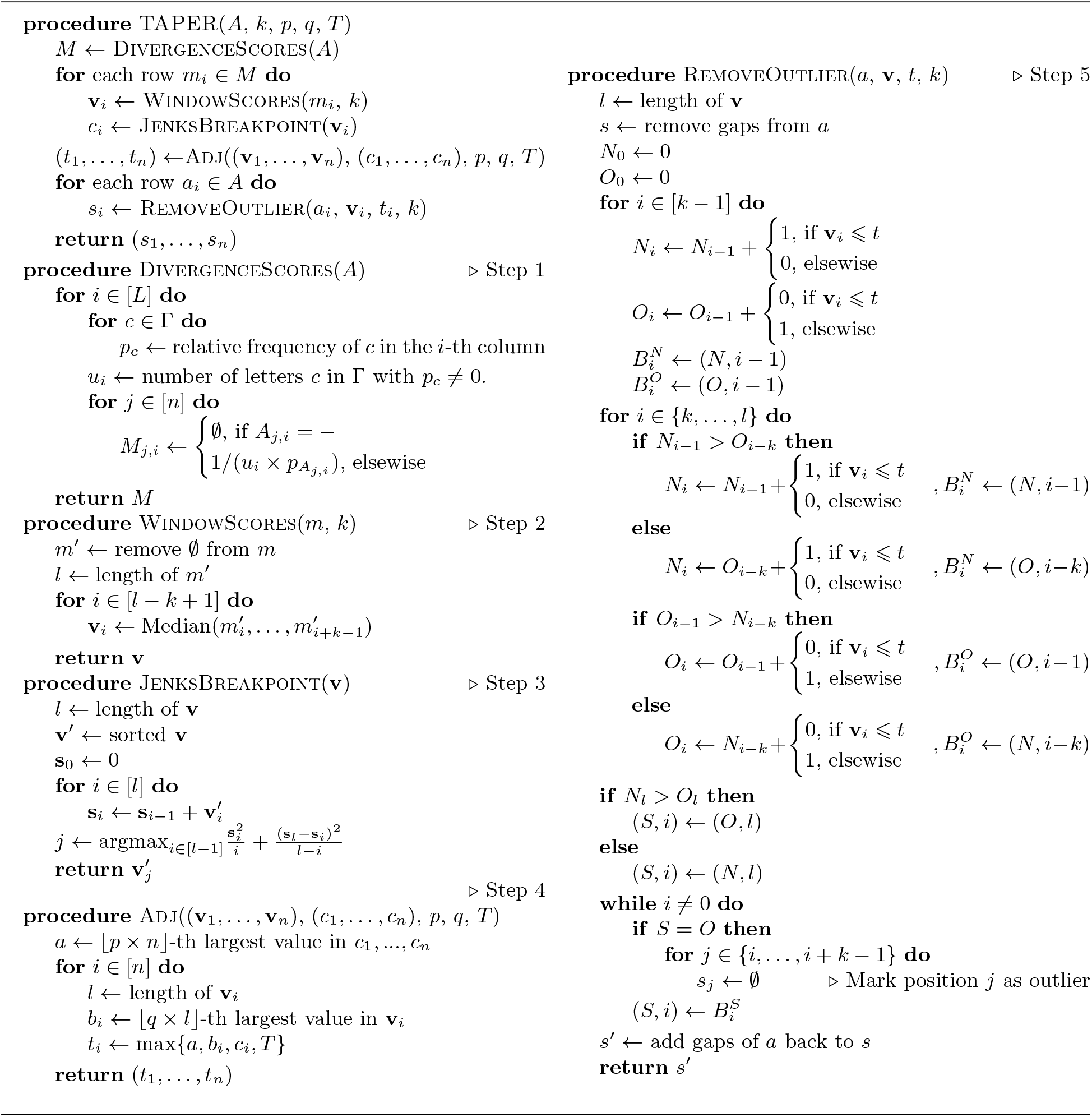
TAPER algorithm. *A*: Input alignment on *n* sequences of length *L* on alphabet Γ and gap letter -. *k, p, q, T*: user-provided parameters. [*X*] denotes 1…*X*.

While several such functions can be imagined, here, we use a scoring method that Henikoff and Henikoff (1992) used for sequence weighting. We score a letter *c* in column *i* as 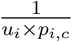 where *u_i_* is the number of unique letters in the column, and *p_i,c_* is the fraction of the letters in the column that are *c*. It can be checked that this score satisfies our criteria (Fig. S1).

#### Step 2

Since per-column scores are noisy, we combine them along small windows *per each sequence*. We first remove gaps from each sequence. Then, selecting an odd constant value *k* (e.g., 11), for each overlapping window of size *k* of each sequence, we assign the median score of the letters in that window as the score of that window. This step produces a distribution of scores for each sequence.

#### Step 3

For each sequence, we seek to find which of the windows have abnormally high scores; these are considered candidate outliers. To do so, we divide windows of *each sequence* into a low scoring and a high scoring group. We find a cutoff point, *t*, such that the squared deviations from the mean of the scores below *t* plus the squared deviations from the mean of points above *t* is minimized. This approach is the Jenks (1967) natural breaks optimization and is equivalent to the 2-means clustering.

#### Step 4

The initial cutoffs from Step 3 include high-scoring windows for all sequences, regardless of whether any outlier exists. To allow sequences with no outliers, we use three parameters 0 < *p, q* < 1 and *T* > 1 to adjust cutoffs. We set the final cutoff of a sequence to the maximum of four quantities: the highest *p*-quantile of all cutoff values across *all* sequences, the highest *q*-quantile of all window scores for *that* sequence, the threshold value *T*, and the initial cutoff *t*. Thus, the adjusted cutoff will include no high-scoring windows for sequences where the windows have homogeneous scores (controlled by *q*) or all scores are within the normal range compared to other sequences (controlled by *p*). User can adjust *p*, *q*, and *T* to control the aggressiveness of the method; *p* controls how many species can have error while *q* controls error length; *T* controls overall aggressiveness. Windows with scores greater than the final cutoff are called red and the rest are called green.

#### Step 5

We divide the original sequence without gaps into alternating *normal* and *outlier* sections. Note that a window can span both sections. The sections boundaries are set using dynamic programming, seeking to maximize the number of red windows fully contained in outlier sections and green windows fully contained in normal sections. The point of this step is smoothing of red and green assignments. Since windows that fall on the section boundaries do not count towards the optimization score, frequent switches between normal and outlier regions are eliminated in this step.

The 5-step procedure we described above is called two-dimensional because scores are computed along the columns (step 1), but outliers are detected (Step 2 and 3) and smoothed (Step 4) along the rows. Thus, for a letter to be marked as an outlier, it must be in several windows, all of which have abnormally high scores compared to the rest of their respective columns, when compared to other windows of the same sequence.

### TAPER Algorithm

2D outlier detection is expected to be more effective at catching smaller errors with smaller values of *k* and longer errors with larger values of *k* (our results confirm this notion; see Fig. S2). To be able to catch a wider range of error lengths, we have devised a strategy to combine several *k* values, each with a different *p, q* setting. We run the 2D outlier algorithm on multiple *k* values and report their union. However, to ensure that a specific *k* value only catches errors of its intended length, we define an upper limit to the length of the detected error, such that only detected errors with lengths less than the upper limit are flagged. In our preliminary analyses, we confirmed that using two or more values of *k* dramatically improves recall for short errors of length 22 (Fig. S3–S4). In our default setting, we use *k* values 5, 9, and 17, with *p* set respectively to 0.25, 0.25, 0.1, and *q* set to 0.1, 0.25, 0.5; we only keep errors of length up to 6 × *k* for *k* = 5 or 9. These settings are motivated by preliminary analyses but are kept fixed as we study various datasets.

## Benchmark

### Evaluation setup

#### Datasets

To benchmark TAPER, we inserted random errors into MSAs of three real datasets with different properties (Table1) and studied whether TAPER can detect them. The 16S.B dataset is an RNA dataset of 16S, with gold-standard alignments built by Cannone *et al.* (2002) and used for benchmarking alignment methods (e.g., Liu *et al.*, 2009; Mirarab *et al.*, 2015; Nguyen *et al.*, 2015). Because this dataset includes 27643 sequences, it enables us to sub-sample the alignment to create subsets with controlled divergence levels. We selected subtrees with a range of diameters (i.e., maximum distance between species) from a phylogeny built from the original MSA. We first find *all* clades in small diameter ranges in increments of 0.025 ([0, 0.025], [0.025, 0.05],…, [0.975, 1]) and then select up to 10 largest clades in each diameter range, requiring at least 20 species. This procedure gave us 371 sub-datasets of the 16S.B dataset, ranging in tree diameter from 0.043 to 0.990. The avian early-bird dataset consists of DNA sequences from 19 genes, aligned automatically but also curated manually by Hackett *et al.* (2008). The RV100-BBA0039 is one of the largest AA alignments available as part of the BAliBASE datasets of gold-standard curated alignments used for benchmarking (Thompson *et al.*, 2005).

**Table 1.** Datasets used in simulations.

| Dataset | Type | # sequences | Seq. Length | Alignment |
| --- | --- | --- | --- | --- |
| 16S.B (371 sub-clades) | RNA | 23–1958 (mean: 544) | 854–2940 (mean: 1362) | structure-based+curated |
| Early-bird (19 genes) | DNA | 72–171 (mean: 158) | 461–2335 (1168) | Mafft + curated |
| Small-AA (RV100-BBA0039) | Protein | 807 | 375 | Gold-standard |

#### Simulating errors

Errors are added to a predefined number of sequences in the alignment (*m*) and for a predefined length (*l*). Sequences with errors are selected uniformly at random. For each of the *m* erroneous sequences, a position to start the error is selected uniformly at random, and *l* of the original non-gap letters are replaced with a randomly chosen letter. For DNA, we choose randomly among the four possible nucleotides. For proteins, we draw the replaced letter from the set of all amino acids such that the chance of selecting each amino acid is proportional to the number of codons that encode to it (e.g., the chance of flipping to Leucine is six times higher than that of Methionine).

Our experiments explore several error profiles (*m* and *l*). First, we fix *m* to be 5% of the total number of sequences, and vary *l* between (2, 3, 5, 8, 16, 32, 64) × 11, including 64 × 11 only for 16S.B as the length is long enough to accommodate such long errors, and excluding 32 × 11 from early-bird genes with mean sequence length below 704 as the error would be more than half of the length. We then fix *l* to 8 × 11 and set *m* to 1 or to 2%, 5%, 10%, or 20% of the number of sequences. To ensure these discrete choices do not impact results, on 16S.B, we also draw *l* and *m* from a normal distribution. We determine *m* by first drawing a value *x* from a normal distribution centered at 20 and a standard deviation of 2 and set 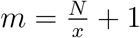, where *N* is the total number of sequences in the alignment (so around 5% of sequences are erroneous). We set the length of the error by drawing from another normal distribution centered at 50 with a standard deviation of 10.

#### Evaluation criteria

We define a False Positive (FP) as any letter that is not an error, but is marked erroneous; a False Negative (FN) as any letter that is erroneous, but is not marked; a True Positive (TP) as any letter that is erroneous, and is marked erroneous; and, a True Negative (TN) as any a letter that is not erroneous, and is not marked. Each letter in the MSA, excluding gap letters, is categorized into one of the four groups, allowing us to compute recall 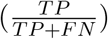 and 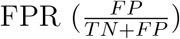. We also report the percentage of the alignment made of errors before 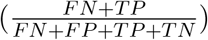 and after 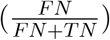 filtering, as well as the percentage of alignment retained 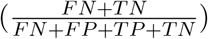.

### Results

#### 16S.B and impact of parameters

On the 16S.B dataset, TAPER is effective in finding erroneous sequences (Fig. 2). The FPR is low, never exceeding 0.16% on average across model conditions, and is even lower before adding errors. Note that we assume that the starting alignment is fully correct, but some FPs may in fact be undetected errors. TAPER retains more than 99% in most cases and never less than 97% of alignment letters (Fig. S5); thus, the method does not overzealously remove data. Depending on the model condition, 70% to 98% of erroneous letters are detected (Fig. 2a). The remaining error is never more than 0.47% of the alignment after filtering, compared to 2.5% before (Fig. 2b).

**Fig. 2.**
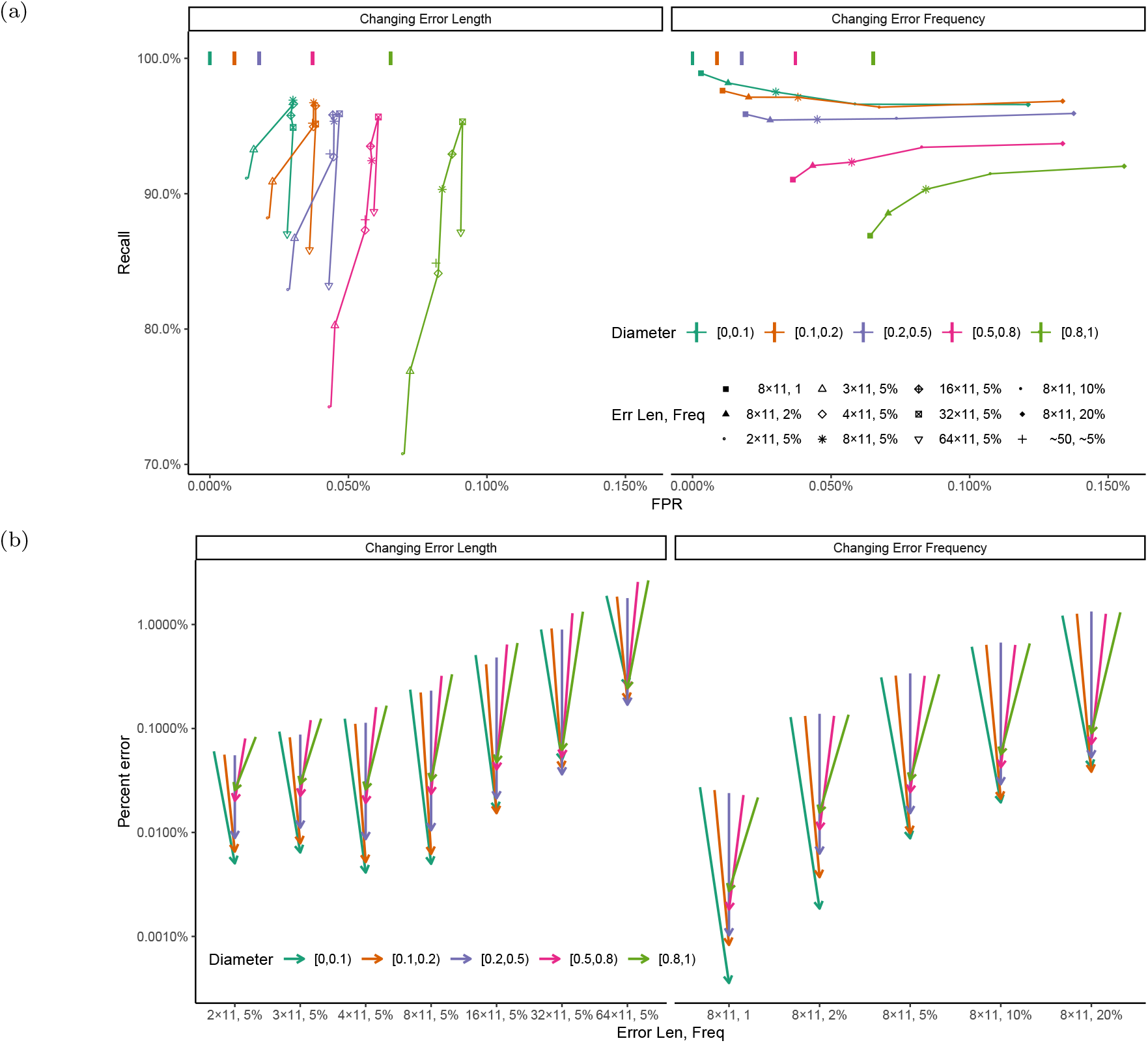
(a) ROC curve (Recall versus FPR) for the subsamples from 16S.B dataset with varying error length, error frequency, grouped into five categories (colours) based on their subtree diameter. Left: Fixing percent of erroneous sequences to 5%, we change the error length from 2 × 11nt to 64 × 11nt. We also draw both length and frequency from Normal distributions, as describe in text (cross). Right: Fixing error length to 8 × 11nt, we vary the number of erroneous sequence between 1 and 2%, 5%, 10%, or 20% of the size of the subtree. Vertical lines on top show the FPR before adding error. See also Figure S6. (b) Reduction in the portion of the alignment nucleotides that are in error. Arrows show percent error before and after filtering. Note the log-scale (see S5b for normal scale).

Tree diameter, which is an indicator of the divergence level, has a substantial impact on the effectiveness of TAPER (Fig. 2 and S6). Over all conditions and using a simple linear model, diameter explains a statistically significant 12% of the variance in the recall (p-value according to an ANOVA test: ≪ 10^−5^; see Table S1). As the diameter increases, recall reduces gradually, but the largest reductions happen when errors are relatively short (Fig. S6). Increasing the diameter also increase FPR, especially when diameter is >0.5. The number of sequences in each alignment has a small impact on the recall (0.5% of total variance) and FPR values (Fig. S7).

FPR is not substantially impacted by the error length but the recall is. Error length explains 32% of variance in the recall (Table S1), which is the lowest with small errors of length 22nt, quickly increases as errors become longer, peaks somewhere between 88 and 352nt, and degrades slightly after that (Fig. 2a). The low recall for short errors should be compared with the amount of error left in the alignment after filtering (Fig. 2b), which is less than 0.03% on average even for high diameters. The amount of remaining error is the highest (0.47% on average) when inserted errors are long, but even then, error has reduced dramatically (2.7% before filtering). When we vary error frequency, we observe small and inconsistent changes in the recall, decreasing for low diameter and increasing for high diameters. FPR increases substantially as the error frequency goes up but remains below 0.16% even when 20% of sequences are erroneous. Finally, when error length and frequency are drawn from a normal distribution, we see consistent results (Fig. 2).

#### Early-bird dataset

On the early-bird dataset, which includes 19 genes, TAPER in most cases has FPR below 0.1% and recall above 60% (Fig. 3a) and reduces the error to less than 1% of the alignment in all cases (Fig. 3b). However, the effectiveness of TAPER varies across genes (Fig. S8a). At one end of the spectrum, on EEF and HMG genes, which have high diameter (1.1 ad 0.97), TAPER has FPR close to 0.1% and relatively low recall (as low as 33% when errors are long and no more than 78% under other conditions). In addition to high diameter, the HMG gene includes 72 out of the 171 species. The other extreme is the BDNF gene (diameter: 0.40) where the mean recall is between 74% and 94% and FPR is below 0.2% across all conditions and below 0.05% in most. While BDNF is known to have patterns of branch length variation that differ from the other genes in the early-bird dataset (cf., Fig. 8 of Braun *et al.*, 2019), several other genes such as NT3 and IRF2 also have high accuracy (Fig. S8). Overall, diameter has modest negative impact on the recall of TAPER and the impact is most noticeable for longer errors (Fig. S8b). According to a linear model only 9% of variation in recall is explained by diameter (p-value: ≪ 10^−5^; see Table S2). For example, the IRF2 gene with a moderately high diameter (0.8) has very high recall (ranging from 71% to 93%). The impact of sequence length and number of sequences on the recall was significant (p-values: 0. < 10^−5^, 0.003) but modest (1.3% and 1% of total variance).

**Fig. 3.**
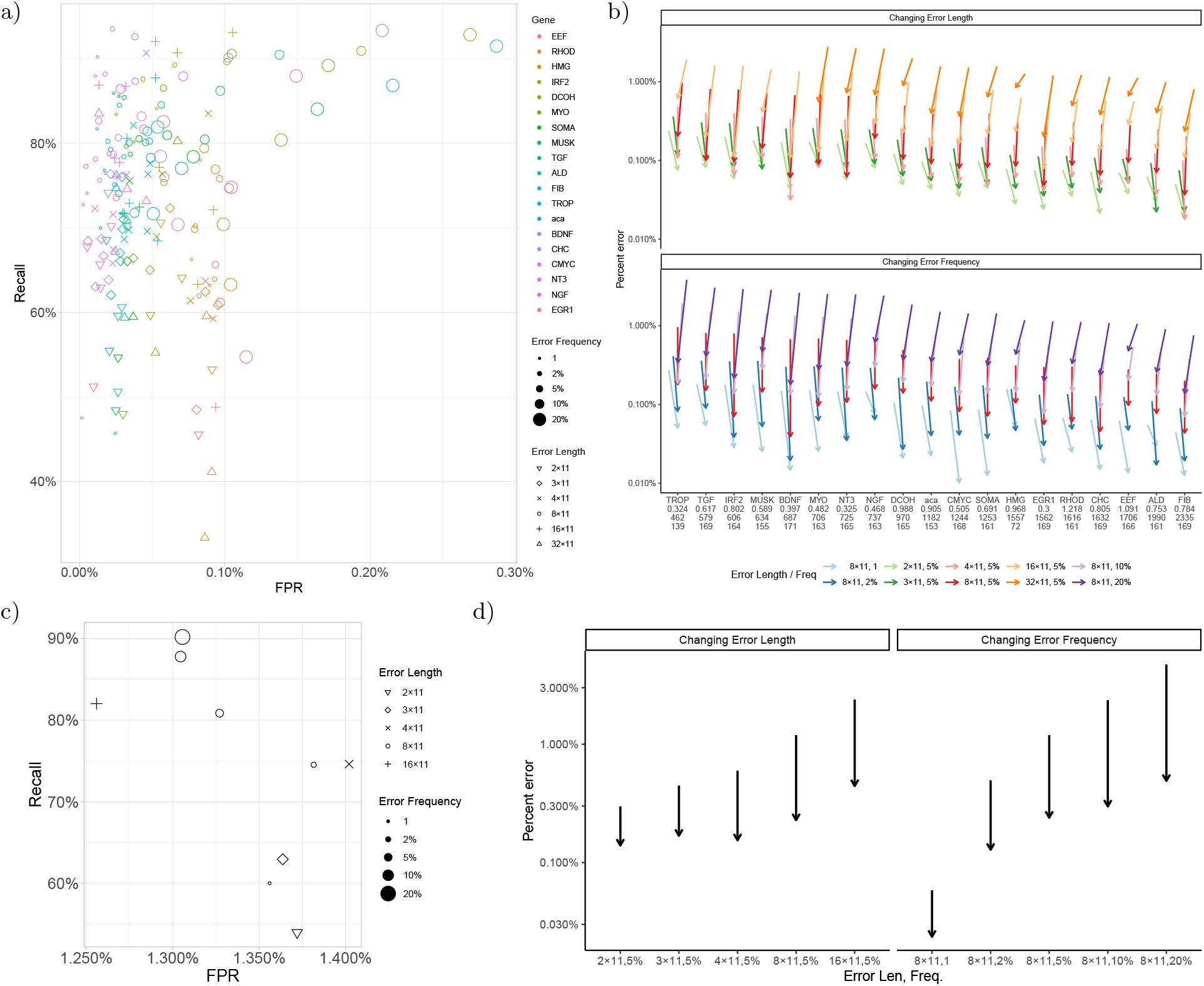
(a,c) ROC results with various error profiles for all genes of the early bird (a) and the AA (c) datasets. (b,d) Reduction in percentage of the nucleotide positions in the alignment that consists of errors for early bird (b) and AA datasets. Arrows show percent error before and after filtering. Note the log-scale. For each gene, below it, we indicate the diameter, mean sequence length, and the number of species present in that gene. See also Fig. S8.

The error profile matters. The error frequency has small impacts on the recall (Table S2), but error length has a large impact (20% of variance; p-value: ≪ 10^−5^). In particular, short errors are difficult for all genes, while long errors are difficult for many but not all genes (e.g., EGR1). Overall, the five factors examined and their interactions explain only 42% of the total variance (Table S2).

#### Small-AA dataset

On the AA dataset, where the original alignment includes a set of sequences that are substantially divergent from the rest (Fig. S10), TAPER has higher FPR (mean: 1.34%) than the previous datasets. We attribute the high FPR to the uncertainty in the alignment, consistent with the observation that even before adding errors, TAPER removes 1.30% of the alignment. This uncertainty makes it harder for TAPER to find inserted errors. Depending on the model condition, the median recall ranges between 54% and 90%. Increasing the frequency of error and the length of the error both improve the recall. A relatively small portion of alignments, typically less than 3%, is removed (Fig. S11) and the remaining error after filtering is never more than 0.5%.

## Biological Examples

To test the effectiveness of TAPER on real data, we revisited 56 genes from the avian dataset of Jarvis *et al.* (2014) analyzed manually by Springer and Gatesy (2018). Springer and Gatesy (2018) found errors in all 56 of these genes (some small and others relatively large). TAPER marked between 0% and 1.2% (mean: 0.2%) of nucleotides in these genes as erroneous (Fig. 4). Since the ground truth of where *all* errors lie is not known, to evaluate results of TAPER, we rely on a likelihood-based metric. If the nucleotides removed are in fact erroneous, we expect the likelihood of the maximum likelihood gene tree to increase, more than it would if we remove the same number of nucleotides from random positions. Further, since gene trees for this dataset are notoriously difficult to estimate accurately (Mirarab *et al.*, 2014), we also compute the likelihood on the best-estimate of the species tree from Zhang *et al.* (2018) (we present this tree as Fig. S12a) after estimating branch lengths for each specific gene; despite a potential for true gene tree discordance, we expect that removing errors would increase the likelihood of the species tree. For both gene trees and the species tree, we see dramatic improvements in the log likelihood (measured using RAxML (Stamatakis, 2014)), far exceeding small increases in the likelihood we get by random removal of the same amount of data (Fig. 4). The more data is removed, the more the likelihood increases. Removing as little as 1% of the data can result in 10% improvements in the log likelihood. Reassuringly, the two outgroups were not removed more often than other species (Fig S12b).

**Fig. 4.**
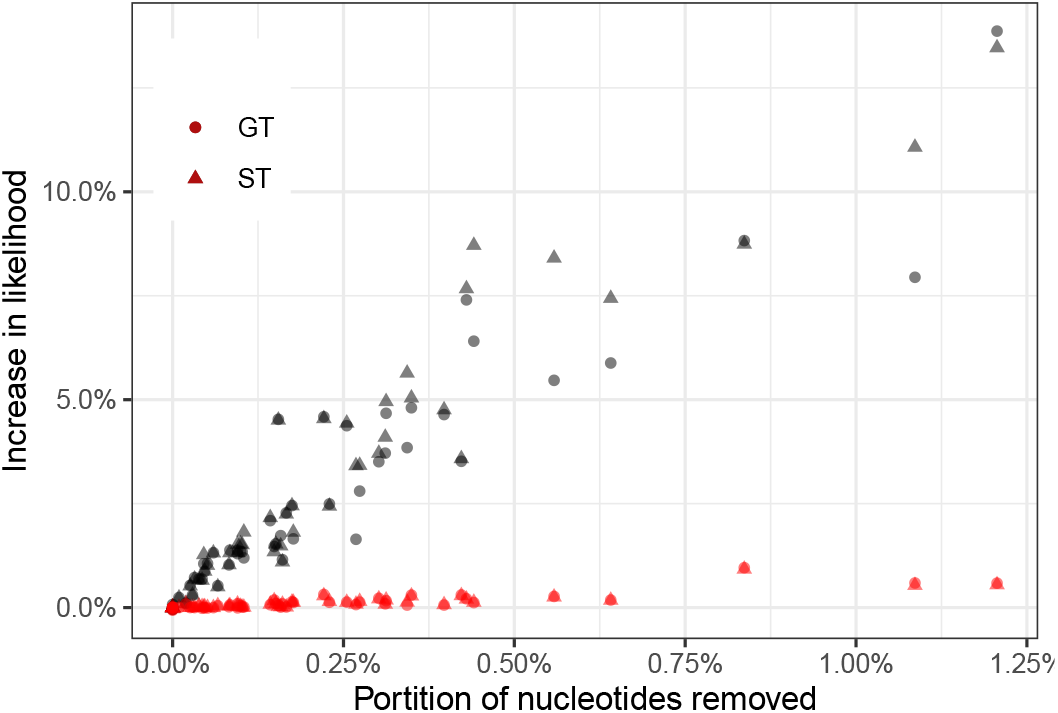
Increase in the log likelihood after filtering computed both for the ML gene tree (GT) and the species tree (ST) versus the portion of the nucleotides in an alignment that are removed. Grey: TAPER filtering. Red: the same amount of data is removed from each gene as TAPER removes but selected from random stretches with the same length. y-axis: change in log-likelihood normalized by likelihood before filtering.

Visual inspection of genes shows that out of 68 cases of error identified by Springer and Gatesy (2018), 24 of them are fully or mostly found by TAPER, and in eight cases, a minority of erroneous positions are found (Table S3). Among the remaining 36 cases (Table S4), 21 are errors that are too short (⩽ 10bp) or too frequent (⩾ 10 out of 48 species) for TAPER to be effective, and in 7 cases, they are somewhat frequent (⩾ 5). In another three cases, only a handful of species are present in those sites, making the errors to be a high *portion* of non-gapped species, which TAPER cannot detect (Table S4). Overall, TAPER marked 21 out of 29 cases that were not too short or too frequent.

### Summary

TAPER was able to reduce error dramatically under varied conditions, but it also had its limitations. TAPER is not very effective in finding very short errors (e.g., below a length 20). On the other extreme, since our method is looking for outlier regions, it cannot detect very large errors (e.g., those that are more than half the sequence length) or those that are too frequent. In between these extremes of short and long is the sweet spot for the error detection by TAPER where there is enough signal to detect oddity of the pattern but the error is not so large that it looks like a real phylogenetic divergence. For shorter errors, changing settings of TAPER (e.g., reducing *k*) could perhaps make the method more sensitive, but that sensitivity would come at the expense of more FP filtering, which we tried hard to avoid. For longer errors, methods like TreeShrink that look for unexpected patterns in branch lengths provide viable alternatives. Besides error length, other factors also mattered. Errors in alignments with very few sequences or very short sequences were harder to detect, as were errors in alignments that were very divergent.

## Disclosure of Potential Conflicts of Interest

The authors declare that there is no conflict of interest.

## Availability

TAPER is implemented fully in Julia and is made available open-source at https://github.com/chaoszhang/AlignmentErrorRemoval. It takes as input a fasta file and outputs a masked alignment.

## Supplementary Materials

## Supplementary figures

**Fig. S1.**
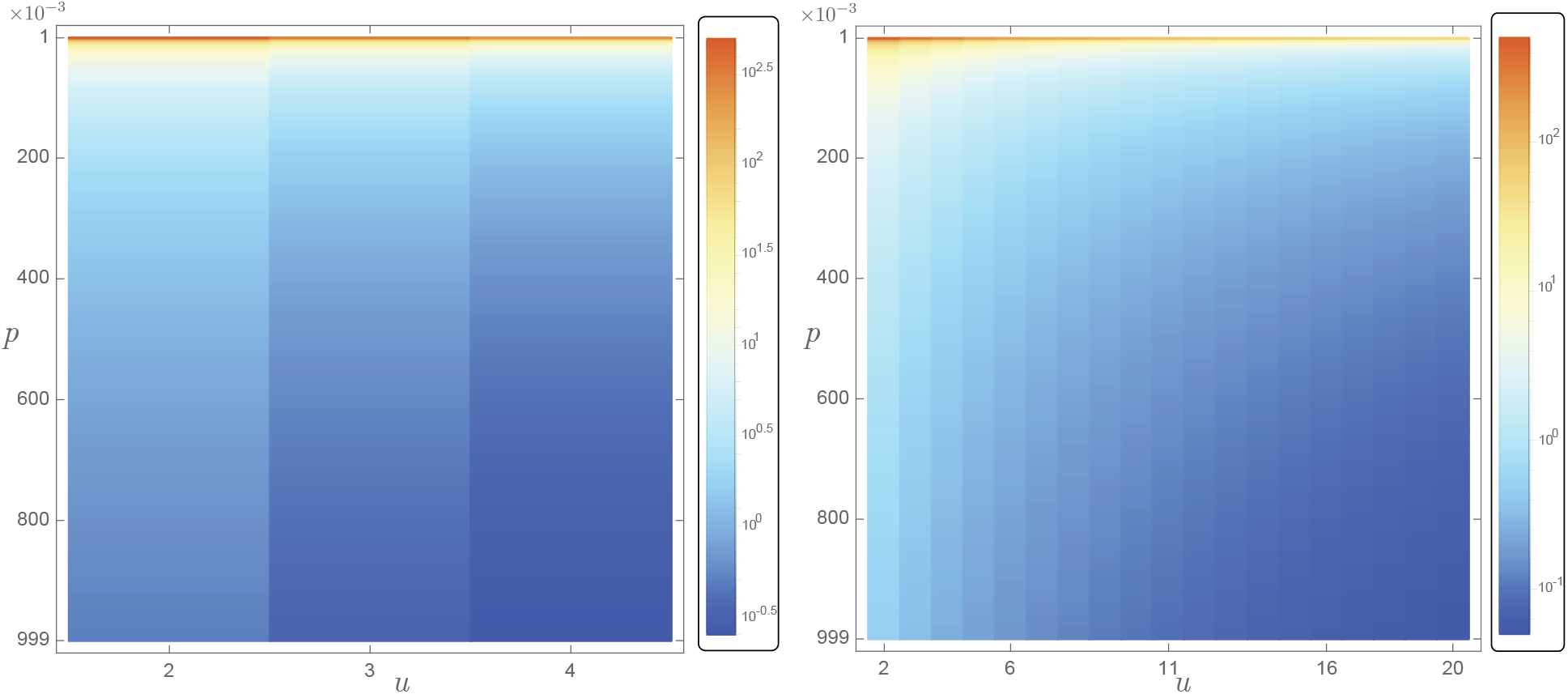
Score function. Left: DNA, Right: Protein.

**Fig. S2.**
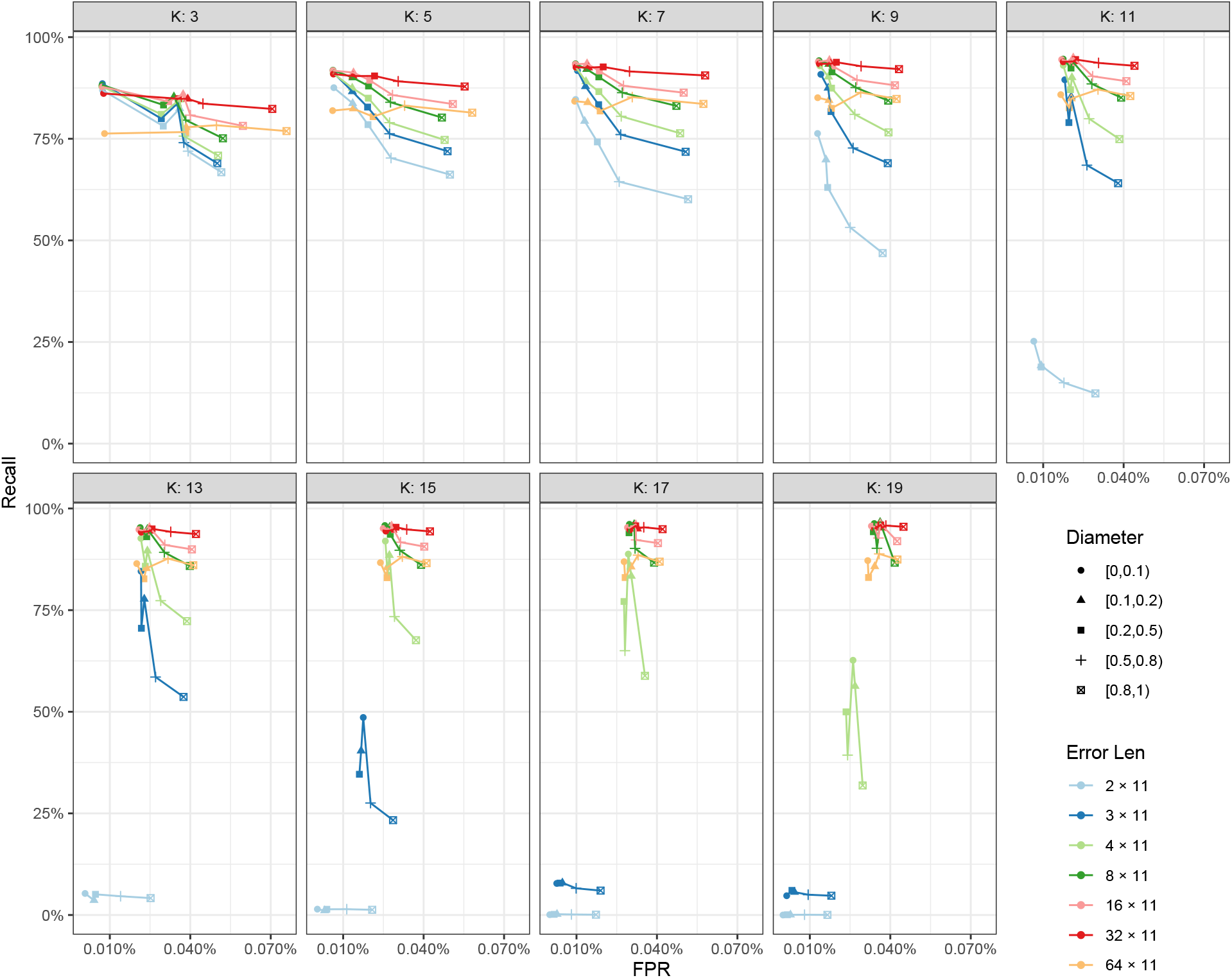
Accuracy of TAPER as we change the parameter *k* (fixing *p* = 0.1 and *q* = 0.5). Smaller *k* is effective for finding shorter errors (e.g., 2 × 11) but less so for finding longer errors (e.g., ⩾ 8 × 11). The false positive rate can increase substantially if *k* is small and errors are long, and the recall is not ideal in those situations. In contrast, larger *k* (e.g. *k* ⩾ 9) is not effective for small errors but can be very effective for longer ones; note how FPR reduces for longer errors when *k* reaches 9.

**Fig. S3.**
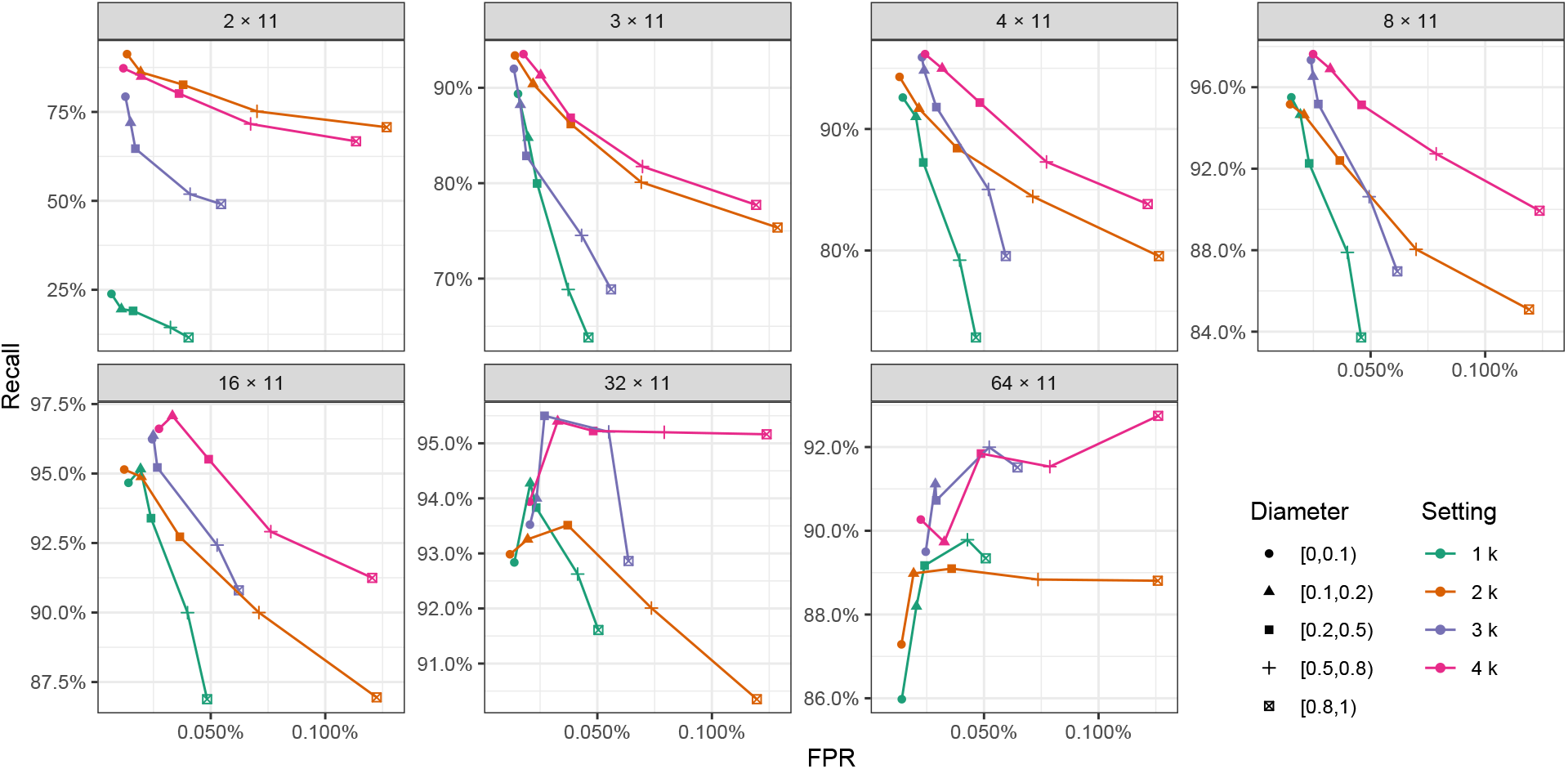
A comparison of various strategies for selecting *k* as the length of error changes (boxes). We either use a single value of *k* = 11 or the union of 2, 3, or 4 values of *k*. When using unions of multiple *k*s, we take results for each *k* only at a certain range of error lengths; for 2k setting: *k* = 5 for error length [0, 30], and *k* = 9 for other lengths; For 3k setting: *k* = 5 for error length [0, 30] *k* = 9 for error length [0, 54] *k* = 19 for other lengths; For 4k setting: *k* = 5 for error length [0, 20] *k* = 7 for error length [0, 35] *k* = 11 for error length [0, 66] *k* = 17 for other lengths. Using one *k* has very low recall in the case with short error length. The other three settings do not universally dominate each other (there is tradeoff between FPR and recall). Overall, the 2k setting seems to have substantially less recall than 4k, with small advantages in FPR. 3k setting generally has better FPR than the other two methods, but also slightly lower recall. Overall, to protect against FP error filtering, we chose the 3k setting that provides a balance between high recall and low FPR. See Figure S4 for more details.

**Fig. S4.**
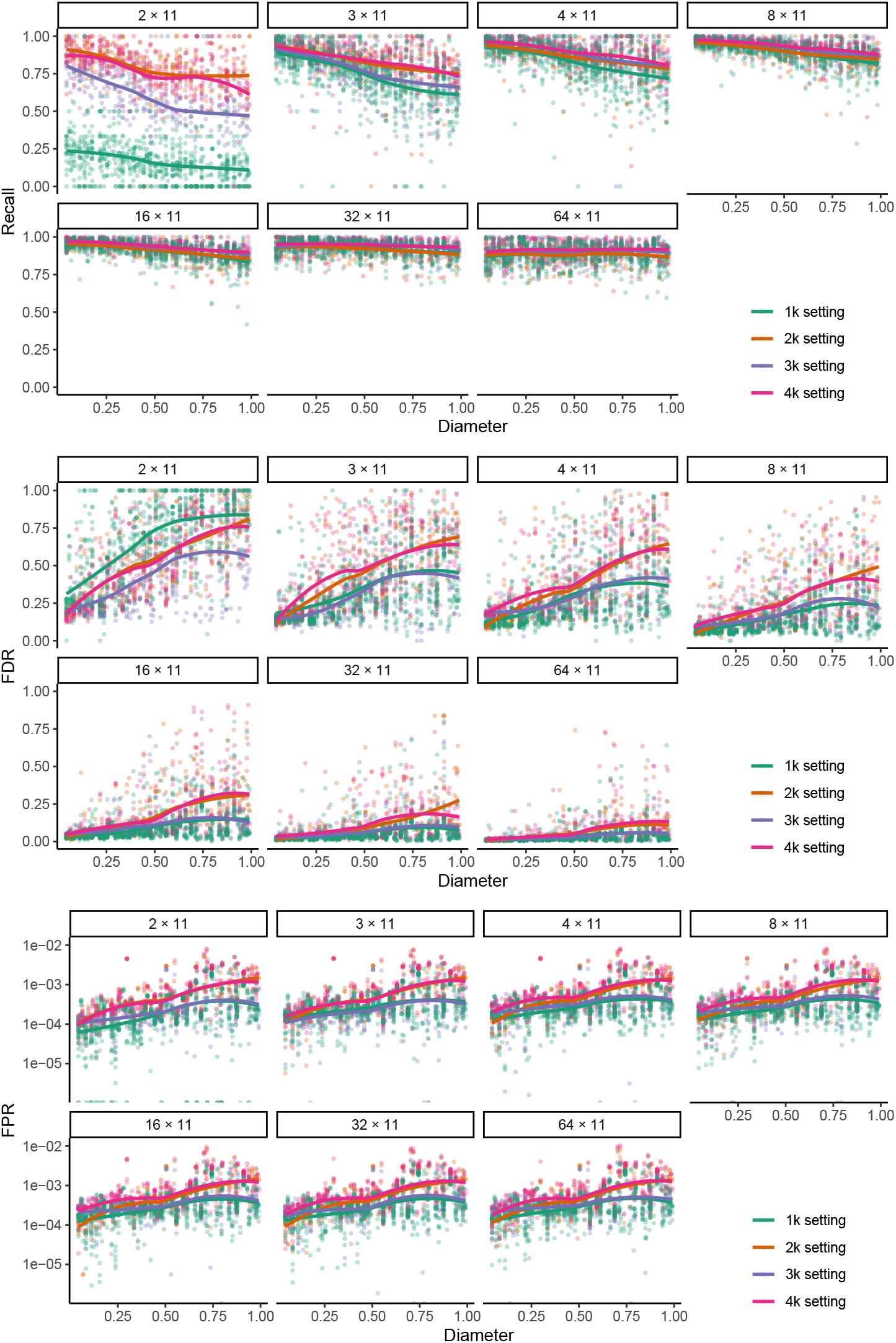
A comparison of various strategies for selecting *k* as the length of error changes (boxes). We either use a single value of *k* = 11 or the union of 2, 3, or 4 values of *k*. When using unions of multiple *k*s, we take results for each *k* only at a certain range of error lengths; for 2k setting: *k* = 5 for error length [0, 30], and *k* = 9 for other lengths; For 3k setting: *k* = 5 for error length [0, 30] *k* = 9 for error length [0, 54] *k* = 17 for other lengths; For 4k setting: *k* = 5 for error length [0, 20] *k* = 7 for error length [0, 35] *k* = 11 for error length [0, 66] *k* = 17 for other lengths. Using one *k* has very low recall in the case with short error length. The other three settings do not universally dominate each other (there is tradeoff between FPR and recall). Overall, the 2k setting seems to have substantially less recall than 4k, with small advantages in FPR. 3k setting generally has better FPR than the other two methods, but also slightly lower recall. Overall, to protect against FP error filtering, we chose the 3k setting that provides a balance between high recall and low FPR.

**Fig. S5.**
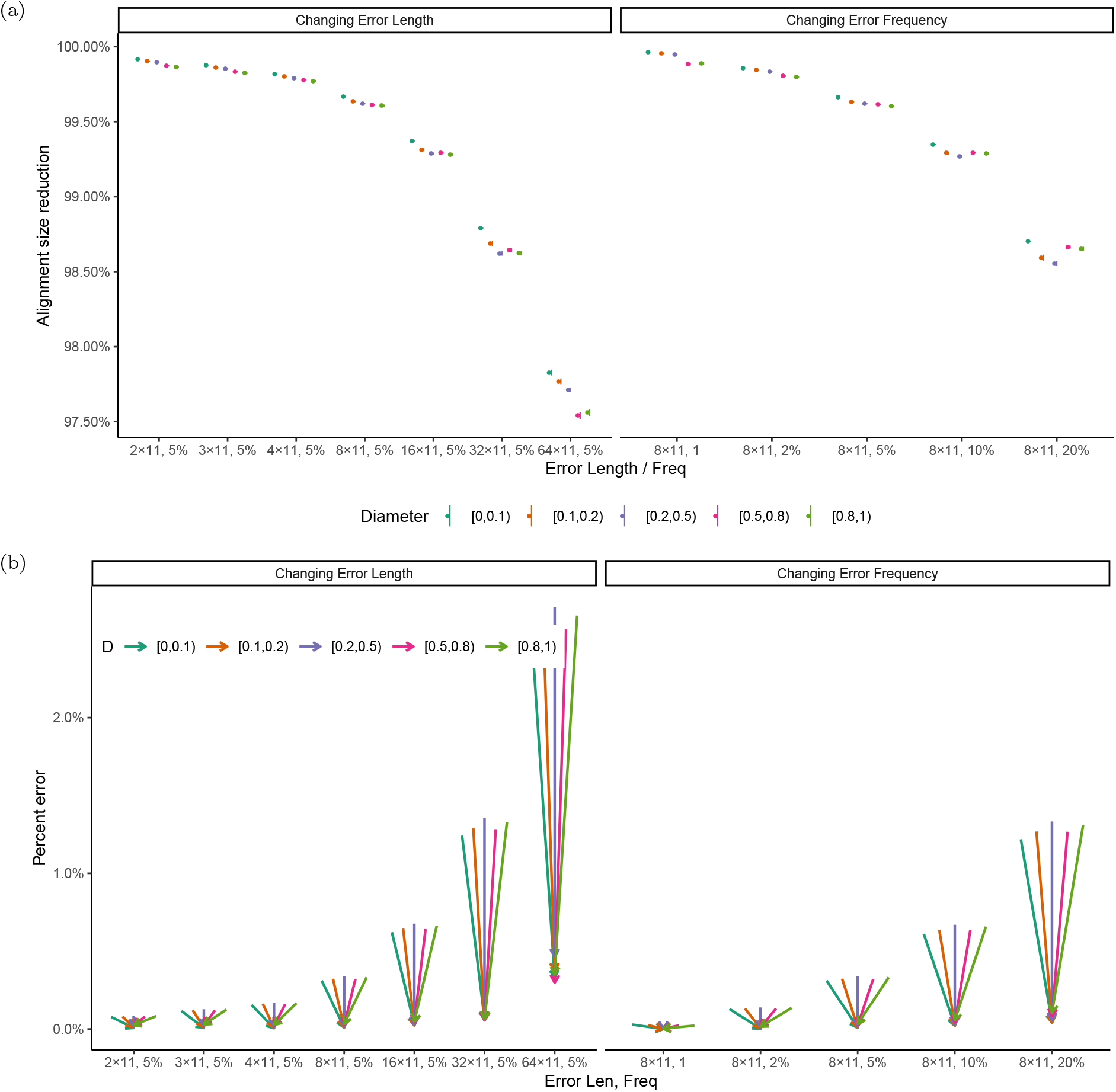
(a) Percentage of the alignment remaining after filtering (the total number of non-gap nucleotide positions in the alignment after divided by before filtering) across model conditions as the error length and frequency changes on the 16S.B dataset. (b) Similar to Figure 2b, we show change in percent error but without log scale.

**Fig. S6.**
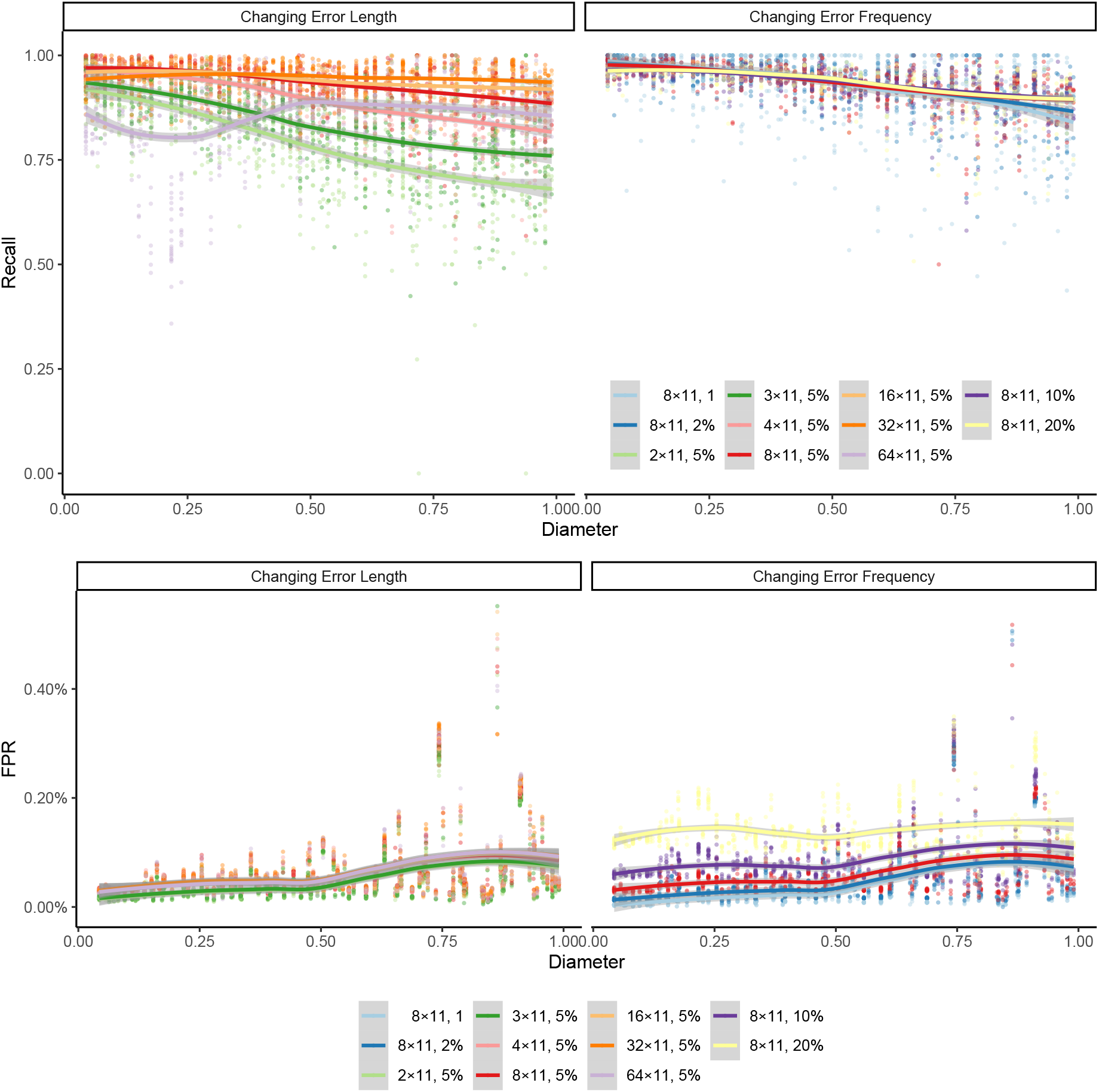
Impact of diameter on Recall and FPR on the 16S dataset.

**Fig. S7.**
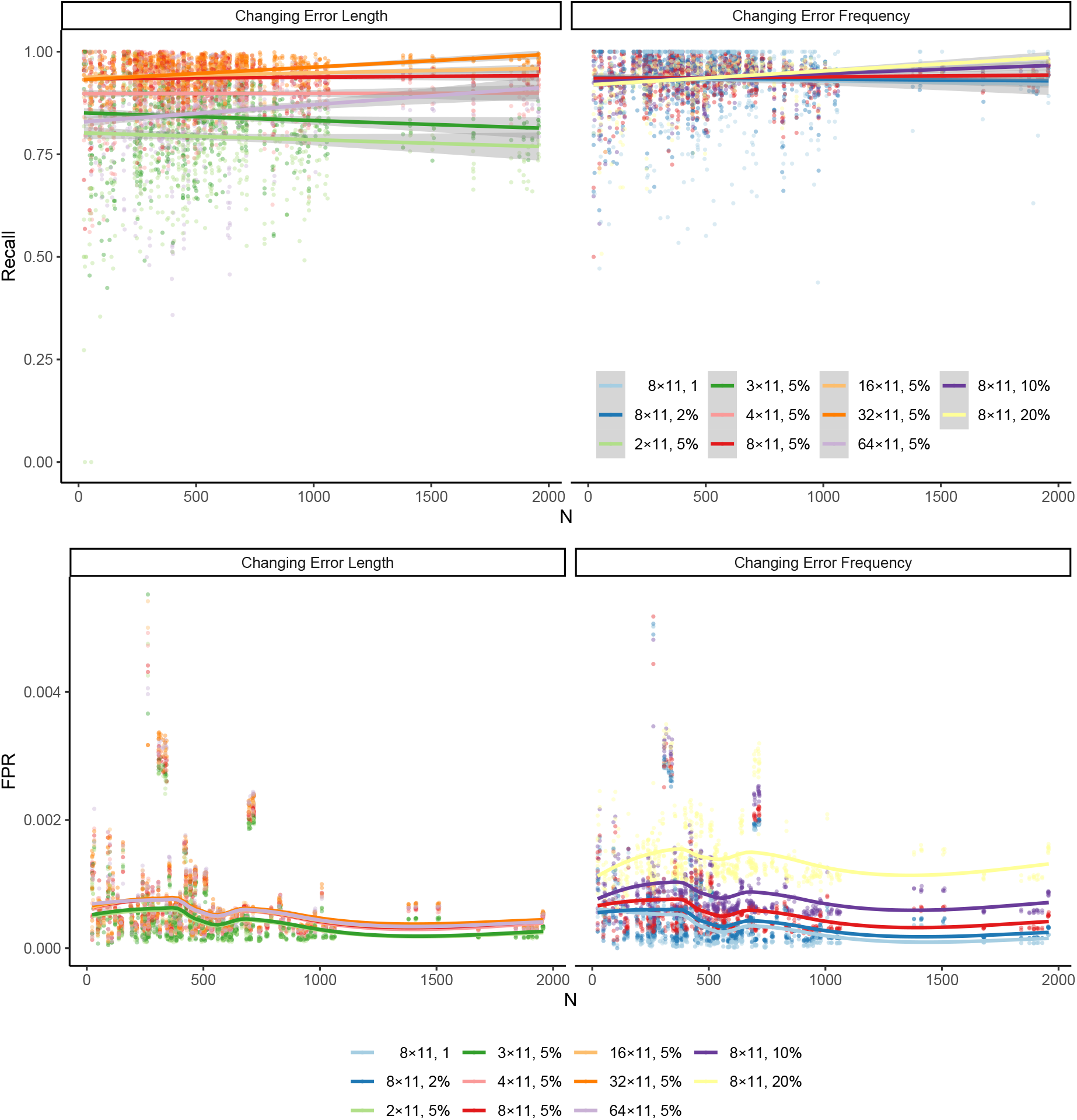
Impact of sequence count on the Recall and FPR on the 16S dataset.

**Fig. S8.**
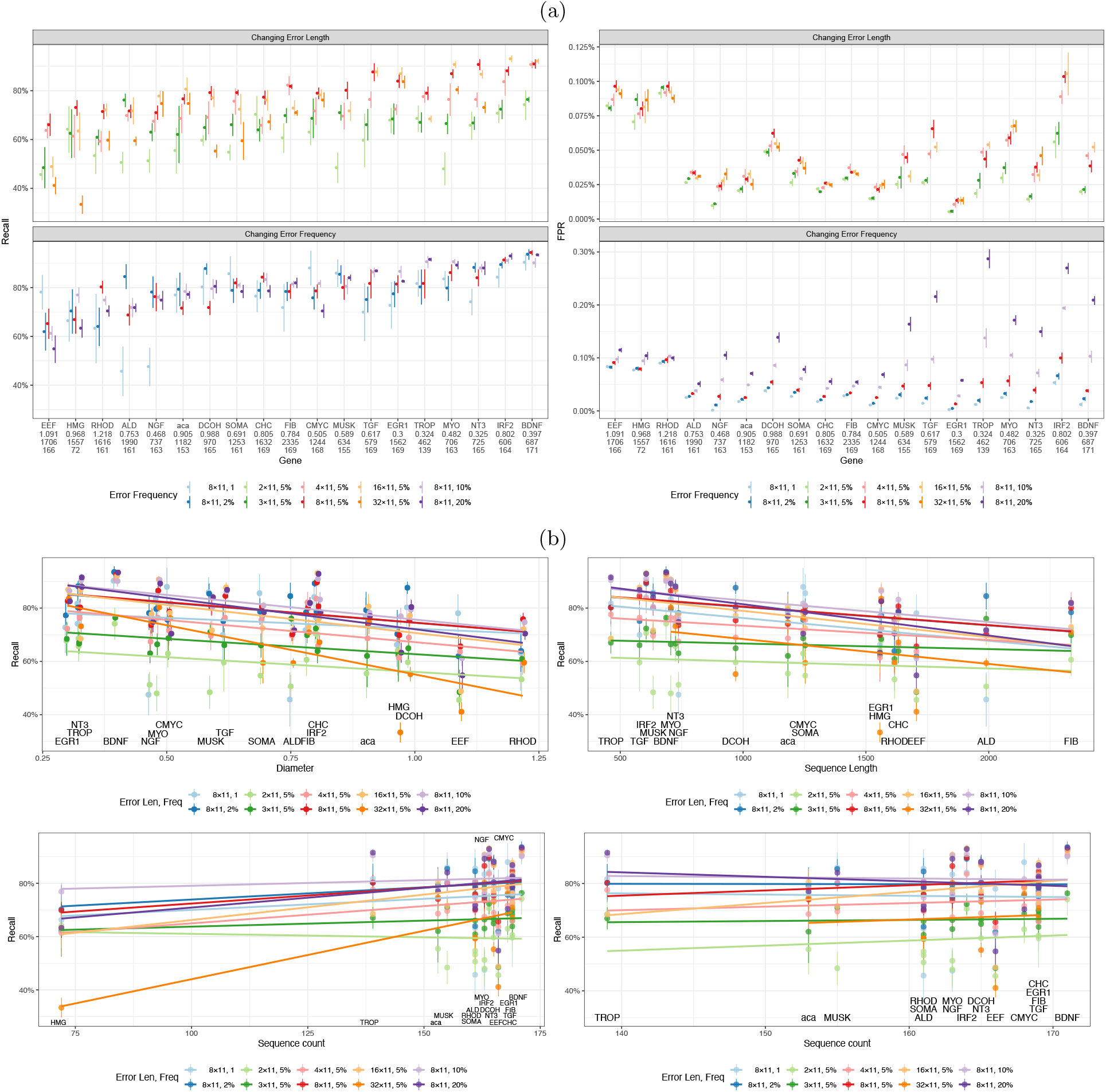
a) Recall and FPR of all genes as we change sequence error or error length. Genes are sorted by their average recall. We show diameter, mean sequence length, and the number of species for each gene under its name. b) Impact of Diameter, sequence count (top left), sequence length (top right), and sequence count (bottom) on the recall. For Sequence count, we show results with and without the outlier gene HMG.

**Fig. S9.**
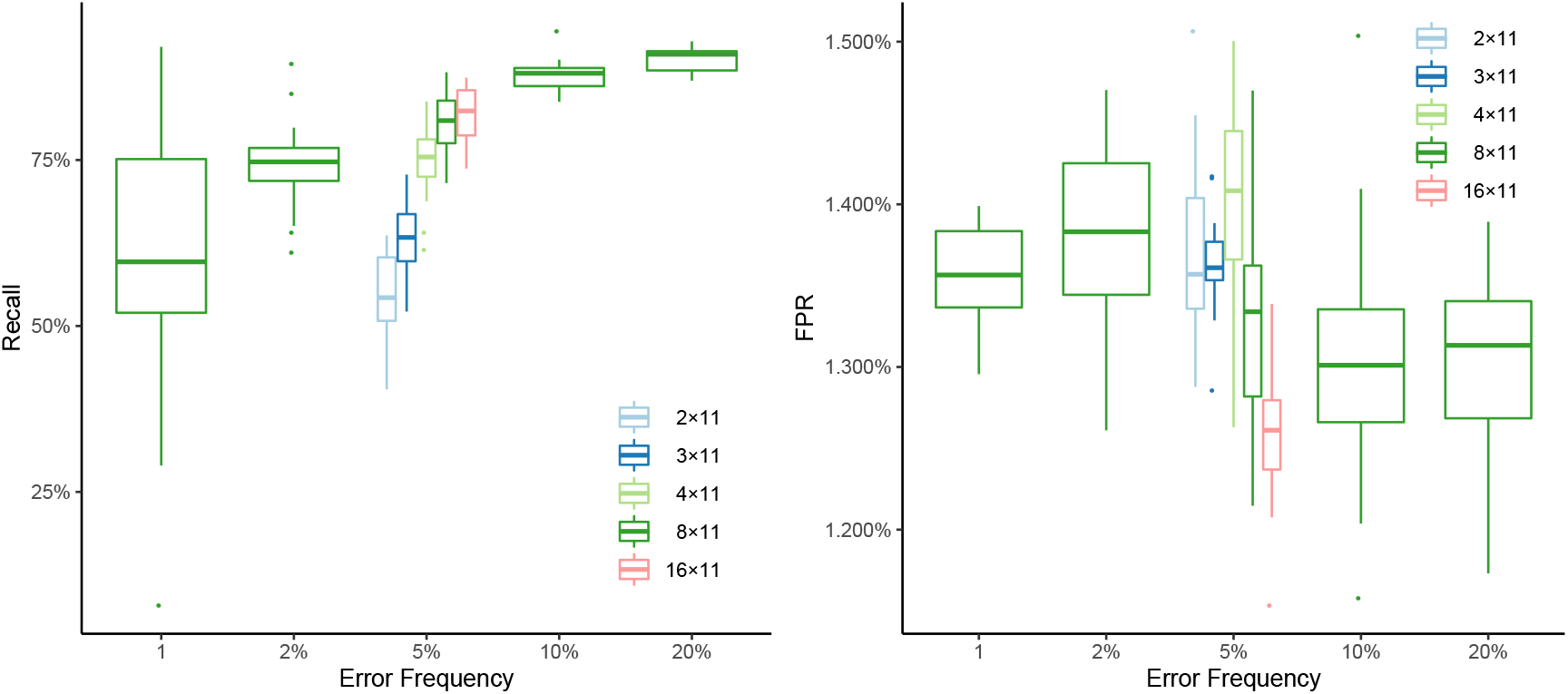
Results on AA dataset.

**Fig. S10.**
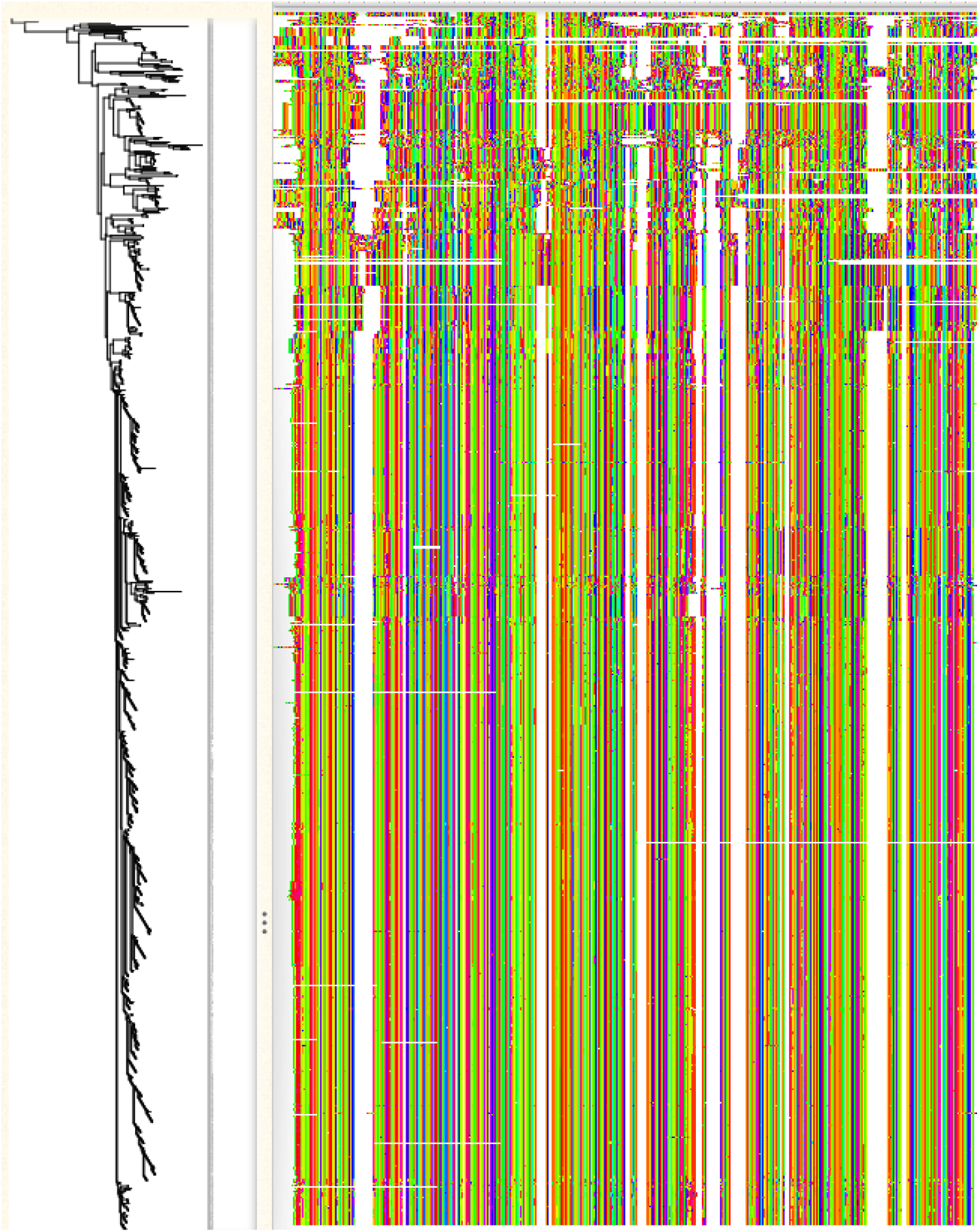
The AA alignment RV100 BBA0039 from the BALIBASE benchmarking dataset. The alignment includes a minority of sites that look very different from the rest.

**Fig. S11.**
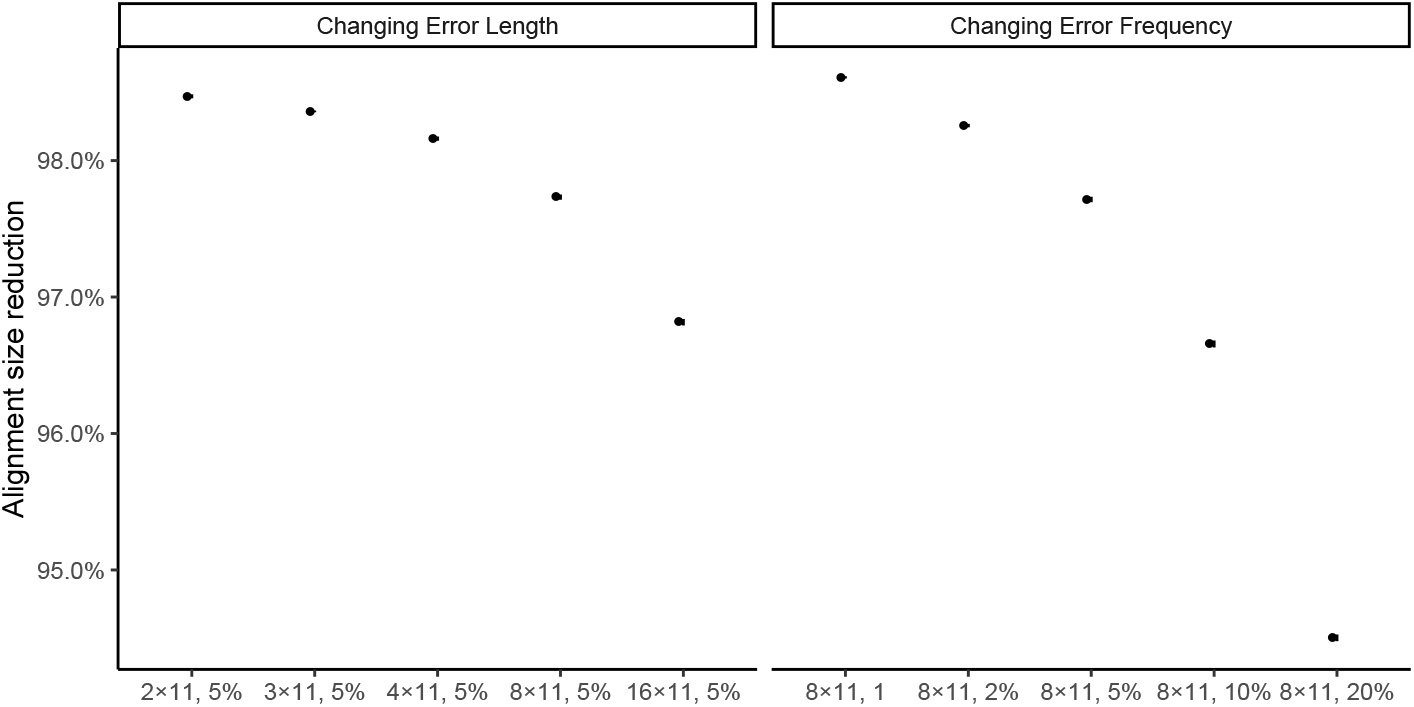
The AA alignment RV100 BBA0039 from the BALIBASE benchmarking dataset. The alignment includes a minority of sites that look very different from the rest.

**Fig. S12.**
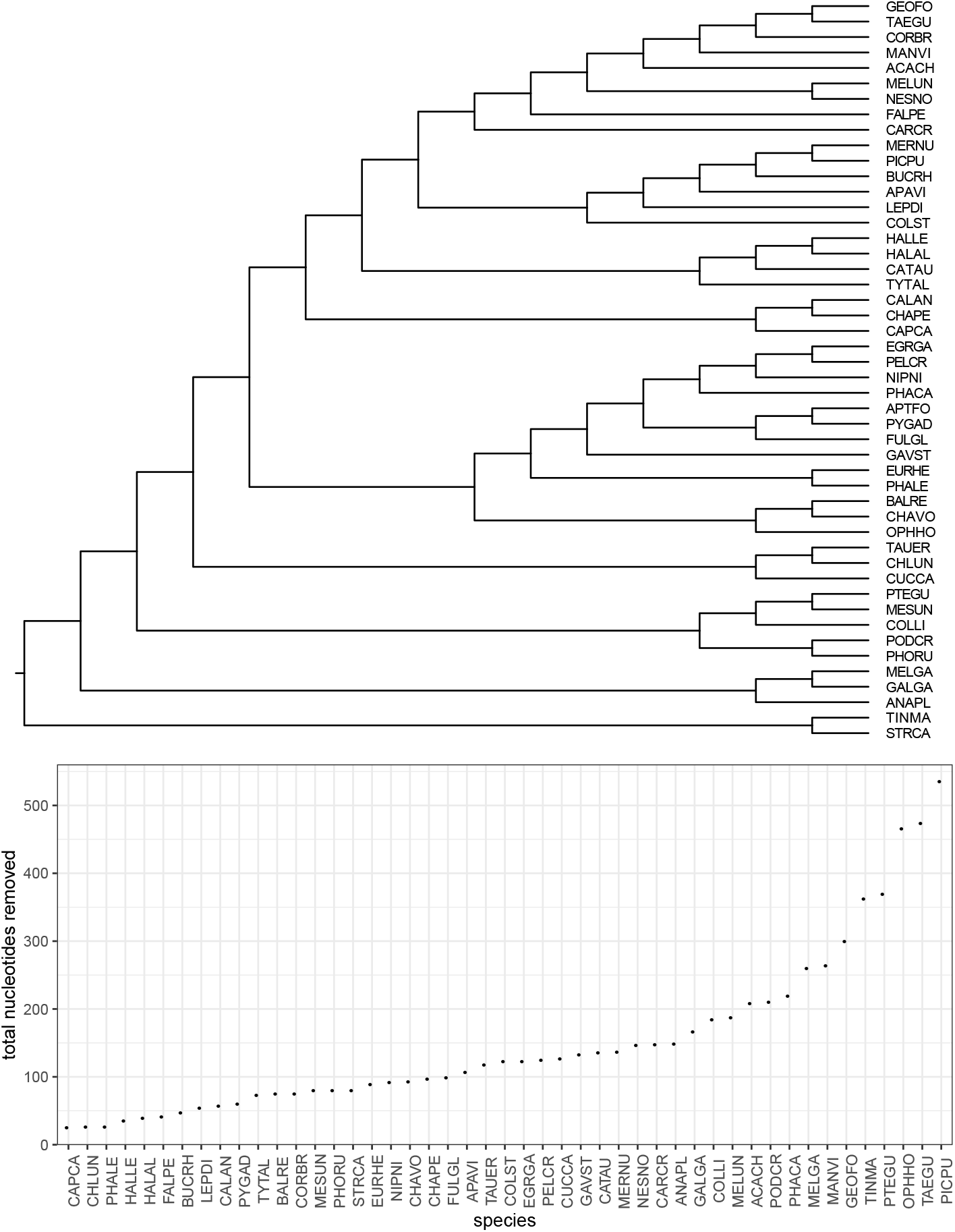
The number of nucleotides removed from species (bottom) does not correspond to phylogenetic relationships (top); in particular, the two outgroups, ostrich (STRCA) and tinamu (TINMA) are not removed more often than others. The species tree shown is obtained using ASTRAL-III run on all > 14,000 input gene trees after contracting branches with support no more than 10%; the tree was reported by Zhang *et al.* (2018).

## Supplementary Tables

**Table S1.** ANOVA test on the 16S dataset, showing impact of four factors and their interactions: Error Length (ErrLen), Error Frequency (n), Diameter, and Sequence Count (N). X:Y corresponds to interactions of variables X and Y.

|  | Df | Sum Sq | Mean Sq | F value | Pvalue | PctExp |
| --- | --- | --- | --- | --- | --- | --- |
| ErrLen | 6 | 21.17 | 3.53 | 991.32 | 0.000000 | 31.9 |
| n | 4 | 0.01 | 0.00 | 0.99 | 0.410483 | 0.0 |
| Diameter | 1 | 7.73 | 7.73 | 2171.08 | 0.000000 | 11.6 |
| N | 1 | 0.32 | 0.32 | 90.54 | 0.000000 | 0.5 |
| ErrLen:Diameter | 6 | 4.94 | 0.82 | 231.22 | 0.000000 | 7.4 |
| n:Diameter | 4 | 0.13 | 0.03 | 9.04 | 0.000000 | 0.2 |
| ErrLen:N | 6 | 0.13 | 0.02 | 6.15 | 0.000002 | 0.2 |
| n:N | 4 | 0.11 | 0.03 | 7.75 | 0.000003 | 0.2 |
| Diameter:N | 1 | 0.03 | 0.03 | 9.44 | 0.002130 | 0.1 |
| ErrLen:Diameter:N | 6 | 0.24 | 0.04 | 11.10 | 0.000000 | 0.4 |
| n:Diameter:N | 4 | 0.00 | 0.00 | 0.17 | 0.951870 | 0.0 |
| Residuals | 8860 | 31.53 | 0.00 |  |  | 47.5 |

**Table S2.**
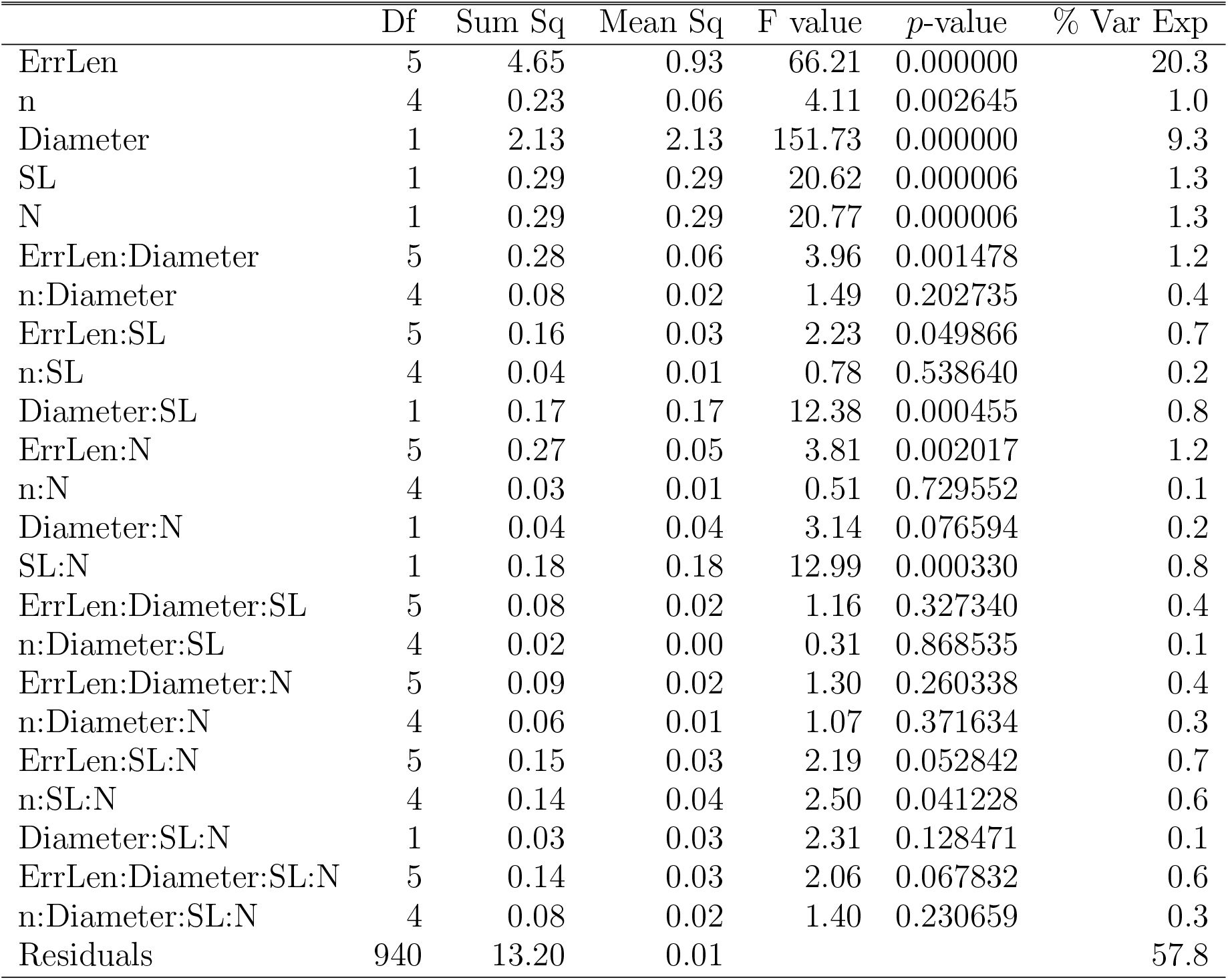
ANOVA test on the early-bird dataset, showing impact of five factors and their interactions: Error Length (ErrLen), Error Frequency (n), Diameter, Sequence Length (SL), and Sequence Count (N). X:Y corresponds to interactions of variables X and Y.

|  | Df | Sum Sq | Mean Sq | F value | *p*-value | % Var Exp |
| --- | --- | --- | --- | --- | --- | --- |
| ErrLen | 5 | 4.65 | 0.93 | 66.21 | 0.000000 | 20.3 |
| n | 4 | 0.23 | 0.06 | 4.11 | 0.002645 | 1.0 |
| Diameter | 1 | 2.13 | 2.13 | 151.73 | 0.000000 | 9.3 |
| SL | 1 | 0.29 | 0.29 | 20.62 | 0.000006 | 1.3 |
| N | 1 | 0.29 | 0.29 | 20.77 | 0.000006 | 1.3 |
| ErrLen:Diameter | 5 | 0.28 | 0.06 | 3.96 | 0.001478 | 1.2 |
| n:Diameter | 4 | 0.08 | 0.02 | 1.49 | 0.202735 | 0.4 |
| ErrLen:SL | 5 | 0.16 | 0.03 | 2.23 | 0.049866 | 0.7 |
| n:SL | 4 | 0.04 | 0.01 | 0.78 | 0.538640 | 0.2 |
| Diameter:SL | 1 | 0.17 | 0.17 | 12.38 | 0.000455 | 0.8 |
| ErrLen:N | 5 | 0.27 | 0.05 | 3.81 | 0.002017 | 1.2 |
| n:N | 4 | 0.03 | 0.01 | 0.51 | 0.729552 | 0.1 |
| Diameter:N | 1 | 0.04 | 0.04 | 3.14 | 0.076594 | 0.2 |
| SL:N | 1 | 0.18 | 0.18 | 12.99 | 0.000330 | 0.8 |
| ErrLen:Diameter:SL | 5 | 0.08 | 0.02 | 1.16 | 0.327340 | 0.4 |
| n:Diameter:SL | 4 | 0.02 | 0.00 | 0.31 | 0.868535 | 0.1 |
| ErrLen:Diameter:N | 5 | 0.09 | 0.02 | 1.30 | 0.260338 | 0.4 |
| n:Diameter:N | 4 | 0.06 | 0.01 | 1.07 | 0.371634 | 0.3 |
| ErrLen:SL:N | 5 | 0.15 | 0.03 | 2.19 | 0.052842 | 0.7 |
| n:SL:N | 4 | 0.14 | 0.04 | 2.50 | 0.041228 | 0.6 |
| Diameter:SL:N | 1 | 0.03 | 0.03 | 2.31 | 0.128471 | 0.1 |
| ErrLen:Diameter:SL:N | 5 | 0.14 | 0.03 | 2.06 | 0.067832 | 0.6 |
| n:Diameter:SL:N | 4 | 0.08 | 0.02 | 1.40 | 0.230659 | 0.3 |
| Residuals | 940 | 13.20 | 0.01 |  |  | 57.8 |

**Table S3.** Errors identified by Springer and Gatesy (2018) that TAPER is able to detect fully (Found), mostly (Majority), or to a lesser degree (Minority). Red: Error is either too short (length ⩽ 10) or involves too many sequences (⩾ 10). Orange: Error involves somewhat high numbers of sequences (between 5 and 10). †: erroneous homology is restricted to a subset of the region identified by Springer and Gatesy (2018). See the supplementary file `supplementary-error-pictures.xlsx` for pictures of errors found.

| Gene | Positions | L | n | Description of error by Springer and Gatesy (2018) | Found |
| --- | --- | --- | --- | --- | --- |
| 1066 | 859-1212 | 353 | 2 | Partial intron 8 (Pterocles, Podiceps) is aligned with different region of intron 8 in Balearica (no homology with other sequences) and exons 9-11 in remaining taxa | Full |
| 82 | 413-498 | 85 | 2 | *Manacus* and *Acanthisitta* exon 6 is combination of intron 5 (5') and exon 6 (3') | Full |
| 1087 | 673-753 | 80 | 2 | Intron 8 (Columba, Anas) aligned against exon 8 in other taxa | Full |
| 1077 | 1150-1206 | 56 | 2 | Intron 11 (Melopsittacus, Colius) aligned with exon 10 in other taxa | Full |
| 1028 | 334-386 | 53 | 2 | *Gavia* and *Struthio* (intron 3) aligned with exon 3 in other taxa | Full |
| 1054 | 577-707 | 30 | 2 | Part of intron 5 (Merops, Nestor) is aligned against exons 6 and 7 in others | Full |
| 1079 | 184-201 | 17 | 2 | 3' end of intron 2 (Eurypyga, Haliaeetus leucocephalus) aligned against exon 2 in other taxa | Full |
| 82 | 136-146 | 11 | 2 | Intron 2 (Cathartes, Tauraco) is aligned with exon 2 in other taxa | Full |
| 16 | 2212-2213 | 2 | 2 | 5 end of intron 21 in Geospiza and Acanthisitta is aligned against 5 end of exon 22 in other taxa | Full |
| 1039 | 1784-1999 | 210 | 3 | Exon 14 sequences in Gavia, Phalacrocorax, and Opisthocomus are poorly aligned with other sequences and show different splice site boundaries | Full |
| 1016 | 82-126 | 44 | 3 | Part of intron 1 and exon 2 (Struthio, Tinamus, Pelecanus) aligned with exon 1 in others | Full |
| 44 | 634-737 | 103 | 4 | Exon 9 in Colius, Acanthisitta, Cariama, and Pterocles is a combination of the 5' region of intron 8 and the 3' region of intron 9 | Full |
| 1042 | 289-310 | 21 | 5 | Exon 1 (Gallus, Haliaeetus leucocephalus, Melopsittacus, Tauraco, Columba) aligned against intron 1 in other taxa | Full |
| 1098 | 161-309 | 140 | 2 | Intron 1 (Opisthocomus, Phalacrocorax) aligned with exon 1 in other taxa | Majority |
| 90 | 733-915 | 80 | 2 | Intron 8 (Charadrius, Tauraco) aligned with exon 9 in other taxa | Majority |
| 1014 | 705-768 | 64 | 2 | Part of intron 7 (Eurypyga and Columba) aligned against exon 8 in others | Majority |
| 1013 | 250-273 | 24 | 2 | *Gallus* and *Tinamus* not homologous with other sequences that are present (which are intron 1); problems with *Meleagris* exon 5 | Majority |
| 93 | 124-139 | 15 | 2 | 3' end of intron 1 (Tyto, Pelecanus, Melopsittacus, Meleagris) is aligned with 3' end of exon 1 in others | Majority |
| 30 | 691-762 | 71 | 3 | Intron 3 (Phoenicopterus, Mesitornis, Leptosomus) aligned with exon 4 of other taxa | Majority |
| 1044 | 342-441 | 99 | 4 | Intron 5 (Opisthocomus, Eurypyga, Balearica, Cathartes) aligned against exon 5 in other taxa | Majority |
| 1039 | 1307-1368 | 61 | 4 | Intron 12 (Corvus, Columba, Pelecanus, Pygoscelis) aligned against exon 12 in others | Majority |
| 89 | 740-796 | 54 | 4 | Four taxa (Gallus, Eurypyga, Fulmarus, Manacus) have partial intron 6 sequence instead of partial exon 6 | Majority |
| 1087 | 1006-1019 | 13 | 5 | 5' end of intron 10 (Cuculus, Apaloderma, Columba, Colius, Anas) aligned against 5' end of exon 10 in others | Majority |
| 1039 | 1360-1524 | 164 | 5? | Exon 11 sequences are poorly aligned across avian tree and have different intron-exon boundaries. The numbers of exons in different taxa also varies. | Majority |
| 1077 | 448-583 | 120 | 2 | Regions of intron 5 (?) and 6 (Melopsittacus, Colius) are aligned with exon 7 in others | Minority |
| 1028 | 1158-1232 | 80 | 2 | Sequences are not homologous in Merops and Nipponia versus other taxa | Minority |
| 1089 | 367-421 | 50 | 2 | Two taxa (Eurypyga, Pygoscelis) have intron 4 aligned against exon 4 in other taxa | Minority |
| 99 | 401-506 | 100 | 7 | Intron 3 (Columba, Pycoides, Chlamydotis, Tauraco, Pygoscelis, Cathartes, Tinamus) aligned against exon 3 in other taxa | Minority |
| 1083 | 127-144 | 17 | 7 | Seven taxa (Chlamydotis, Anas, Tinamus, Cariama, Tauraco, Acanthisitta, Opisthocomus) with intron 2 aligned against exon 2 in others | Minority |
| 1097 | 912-1120 | 200 | 10 | Intron 6 in some taxa (e.g., Tinamus) aligned with exon 6 in other taxa (e.g., Struthio) | Minority |
| 28 | 1-81† (48-80) | 32 | 13 | Intron 2 in 13 taxa (e.g., Haliaeetus albicilla) is aligned with exon 1 and exon 2 in others (e.g., H. leucocephalus) | Minority |
| 20 | 754-842 | 88 | 14 | 5' region of intron 1 (Gallus and others) aligned against 3' region of intron 1 (e.g., Falco and others); middle portion of intron 1 (Gallus group) aligned against 5' end of exon 2 (Falco group). Also some taxa in Gallus group (palaeognaths, Cathartes, Podiceps, Charadrius) are misaligned with others in group | Minority |

**Table S4.**
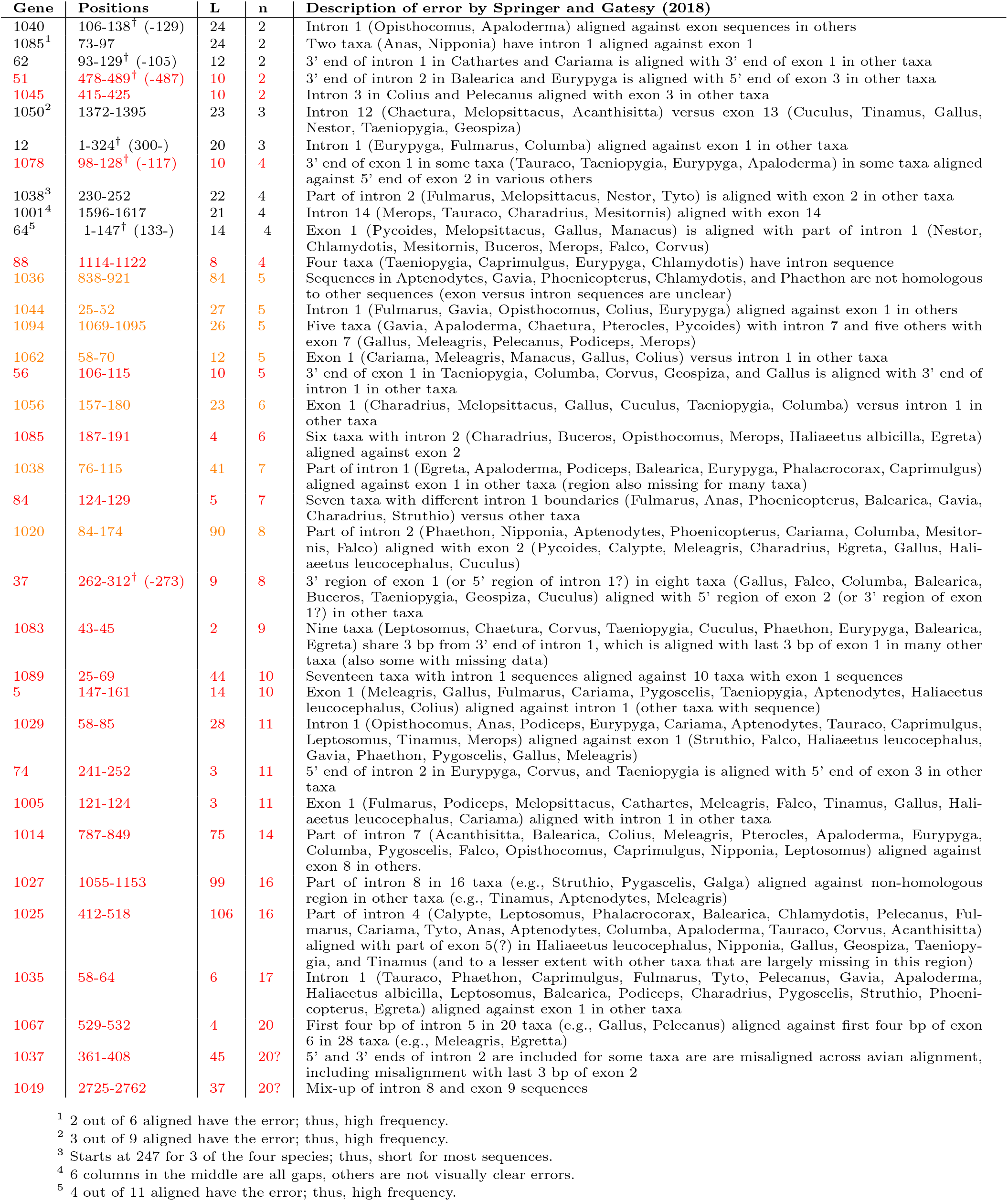
Errors identified by Springer and Gatesy (2018) and missed by TAPER. Notations same as Table S3. †: real error boundary. Error is short (⩽ 10bp) or frequent (⩾ 10); Error is somewhat frequent (⩾ 5, ⩽ 10).

## Notes

### Competing Interest Statement

The authors have declared no competing interest.

